# On the evolution of chaperones and co-chaperones and the expansion of proteomes across the Tree of Life

**DOI:** 10.1101/2020.06.08.140319

**Authors:** Mathieu E. Rebeaud, Saurav Mallik, Pierre Goloubinoff, Dan S. Tawfik

**Author notes:** Equally contributing first authors (Mathieu E. Rebeaud, Saurav Mallik). Lead Contact: D.S.T.

## Abstract

Across the Tree of Life (ToL), the complexity of proteomes varies widely. Our systematic analysis depicts that from the simplest archaea to mammals, the total number of proteins per proteome expanded ~200-fold. Individual proteins also became larger, and multi-domain proteins expanded ~50-fold. Apart from duplication and divergence of existing proteins, completely new proteins were born. Along the ToL, the number of different folds expanded ~5-fold and fold-combinations ~20-fold. Proteins prone to misfolding and aggregation, such as repeat and beta-rich proteins, proliferated ~600-fold, and accordingly, proteins predicted as aggregation-prone became 6-fold more frequent in mammalian compared to bacterial proteomes. To control the quality of these expanding proteomes, core-chaperones, ranging from HSP20s that prevent aggregation to HSP60, HSP70, HSP90, and HSP100 acting as ATP-fueled unfolding and refolding machines, also evolved. However, these core-chaperones were already available in prokaryotes, and they comprise ~0.3% of all genes from archaea to mammals. This challenge—roughly the same number of core-chaperones supporting a massive expansion of proteomes, was met by (i) higher cellular abundances of the ancient generalist core-chaperones, and (ii) continuous emergence of new substrate-binding and nucleotide-exchange factor co-chaperones that function cooperatively with core-chaperones, as a network.

## Introduction

All cellular life is thought to have stemmed from the Last Universal Common Ancestor, LUCA (Doolittle, 1999; Woese, 1998), that emerged more than 3.6 billion years ago. Two major kingdoms of life diverged from LUCA: bacteria and archaea, which about 2 billion years later merged into the eukaryotes (Eme et al., 2017). Since the beginning of biological evolution, life’s volume has increased on a grand scale: the average size of individual cells has increased ~100-fold from prokaryotes to eukaryotes (Cooper and Hausman, 2000), the number of cell types has increased ~200-fold from unicellular eukaryotes to humans (Arendt, 2008), and average body size increased ~5000 fold from simplest sponges to blue whales (Heim et al., 2017).

This expansion in organismal complexity and variability was accompanied by an expansion in life’s molecular workforce, proteomes in particular, which in turn presented a challenge of reaching and maintaining properly folded and functional proteomes. Most proteins must fold to their native structure in order to function, and their folding is largely imprinted in their primary amino acid sequence (Anfinsen, 1973; Dill and Chan, 1997; Englander and Mayne, 2014). However, many proteins, and especially large multi-domain polypeptides, or certain protein types such as all-beta or repeat proteins, tend to misfold and aggregate into inactive species that may also be toxic (Han et al., 2007). Life met this challenge by evolving molecular chaperones that can minimize protein misfolding and aggregation, even under stressful out-of-equilibrium conditions favoring aggregation (Finka et al., 2016; Kim et al., 2013). Chaperones can be broadly divided into core- and co-chaperones. Core-chaperones can function on their own, and include ATPases HSP60, HSP70, HSP100, and HSP90 and the ATP-independent HSP20. The basal protein holding, unfolding, and refolding activities of the core-chaperones are facilitated and modulated by a range of co-chaperones such as J-domain proteins (Caplan, 2003; Duncan et al., 2015; Schopf et al., 2017).

Starting from LUCA, as proteomes expanded, so did the core-chaperones and their respective co-chaperones. Indeed, chaperones have been shown to facilitate the acquisition of destabilizing mutations and thereby accelerate protein evolution (Alvarez-Ponce et al., 2019; Bogumil and Dagan, 2012; Tokuriki and Tawfik, 2009). However, the co-expansion of proteomes and of chaperones, underscoring a critical balance between evolutionary innovation and foldability, remains largely unexplored. We thus embarked on a systematic bioinformatics analysis that explores the evolution of both proteome and chaperones, and of both core- and their auxiliary co-chaperones, along the tree of life.

## Results

### A Tree of Life analysis of the expansion of proteomes and chaperones

We aimed to explore, systematically, across the Tree of Life (ToL) the expansion of proteomes and compare it to the chaperone composition and level. To this end, we collected proteome sequences from representative organisms belonging to all the major bacterial, archaeal, and eukaryotic clades and constructed a ToL (**Figure 1A**; **Data S1**; see also **Methods**). The overall topology of our tree was borrowed from TimeTree (Hedges et al., 2015) and we also adhered to their order of divergence which is based on molecular dating and geological records. TimeTree also provides putative dates of emergence and these are provided as branch lengths, yet because our analysis is primarily comparative, branch lengths were only used here as graphical aid (**Figure 1A**).

**Figure 1.**
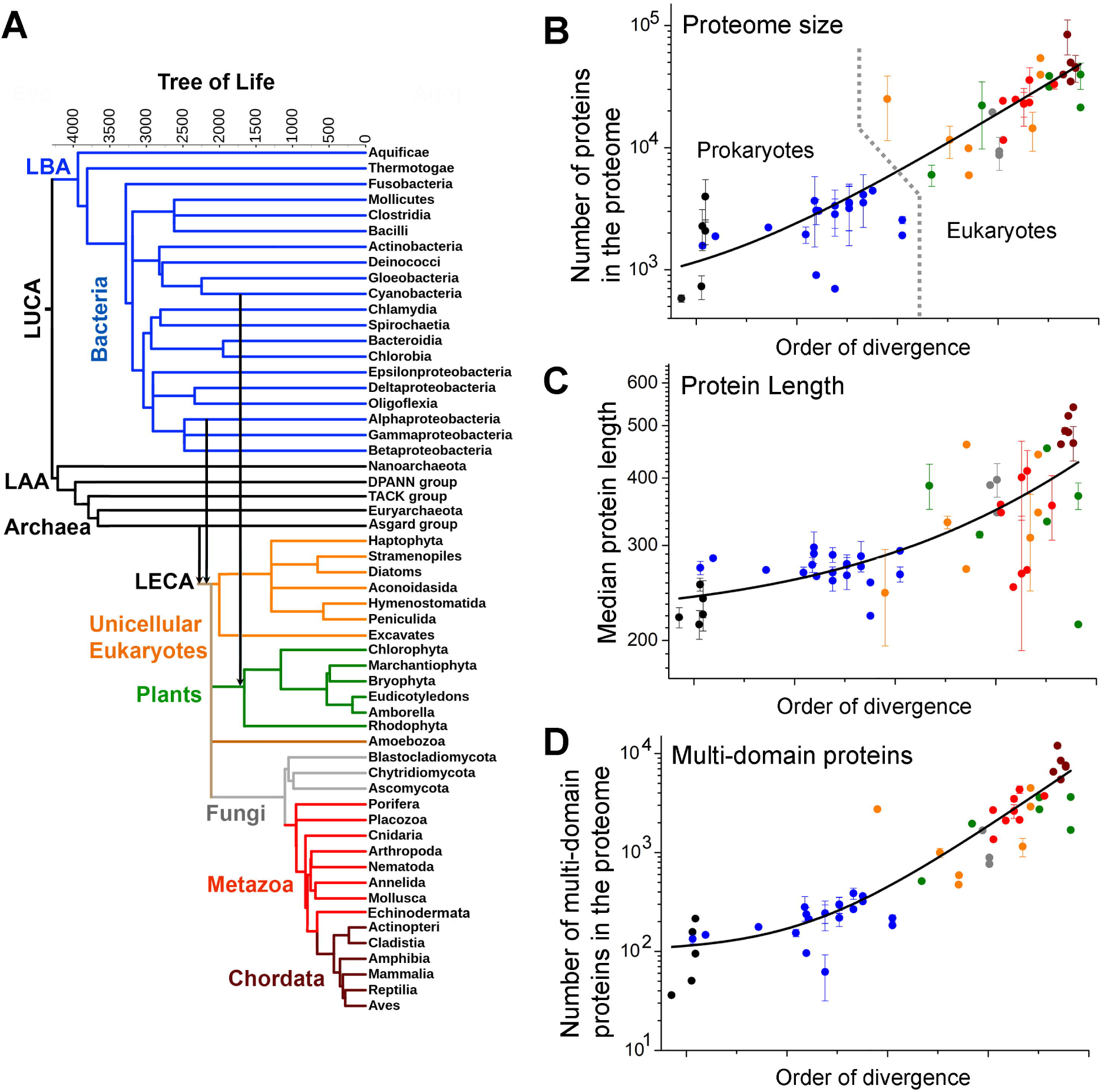
Expansion of proteome size across the Tree of Life. **(A)** The Tree of Life (ToL) used in this study. Leaves represent extant phylogenetic clades, while internal nodes represent their presumed ancestors. Branch lengths are in million years, as available from TimeTree (Hedges et al., 2015), and they refer to the relative order of divergence of the corresponding clades rather than the absolute dates of their emergence. The major phylogenetic groups in the tree (Bacteria, Archaea, Unicellular Eukaryotes, Plants, Fungi, Metazoan, and Chordata) are highlighted in different colors. Vertical arrows highlight the two major endosymbiosis events: the Alphaproteobacterial origin of mitochondria and the Cyanobacterial origin of plastids. Abbreviations: LUCA: Last Universal Common Ancestor, LAA: Last Archaeal Ancestor, LBA: Last Bacterial Ancestor, LECA: Last Eukaryotic Common Ancestor. **(B**□**D)** The average per clade values for various proteome size parameters (Y-axis, in log scale) are plotted against their order of divergence (along X-axis in linear scale). These parameters include proteome size **(B)**, median protein length, **(C)**, and multi-domains proteins in the proteome **(D)**. In these scatterplots, the colors of data-points represent their major phylogenetic group in the tree (panel **A**). Error bars represent the clade standard deviation (no error bars relate to clades comprising only one representative organism). The lines were derived by a fit to an exponential equation, and are provided merely as visual guides. Prokaryotic and eukaryotic organisms were separated by a dashed line in panel-B.

The Tree of Life begins with the Last Universal Common Ancestor (LUCA) at the root (**Figure 1A**). The edges of the ToL represent the extant three kingdoms – archaea, plotted throughout in black, bacteria, plotted in blue, and eukaryotes. The latter emerged by endosymbiosis of an *Alphaproteobacterium* and an Asgard-like archaeon (de Duve, 2007; Margulis et al., 2006). The emergence of green algae and subsequently of plants occurred with a secondary endosymbiosis of a *Cyanobacterium* into a non-photosynthetic eukaryote (Martin et al., 1998). The major eukaryotic clades therefore comprised: unicellular, early diverging eukaryotes (in orange), fungi (grey), plants (green), metazoa (invertebrate animals, red), and Chordata (vertebrate animals; wine). Overall, our analysis was based on comparing the proteomes of 188 representative organisms, covering 56 major clades of bacteria, archaea, and eukaryote (**Data S1**). The various proteome parameters analyzed below were initially derived for each representative organism in the core-tree. The representative organisms of each clade were then pulled together to calculate the clade average and the standard deviation for this average. The clade average values were subsequently plotted using the order of divergence for the X-axis. Accordingly, these plots also broadly divide to prokaryotes (the left part) and eukaryotes (the right part), and the latter’s right edge comprises chordata including Mammalia (**Figure 1B**, and the following figures).

### The expansion of proteome size

Initially, we scrutinized the expansion of proteome size by examining (i) the total number of proteins per proteome in a given clade; (ii) the median protein length; and (iii) the number of multi-domain proteins in the proteome. The clade average values of these three parameters were plotted in **Figure 1B□D**, with colors of points matching with the branch-colors in **Figure 1A**).

#### *The total number of proteins per proteome expanded ~200-fold* (Figure 1B)

Proteomes that comprise a larger number of proteins unavoidably present a greater challenge for their protein quality control chaperone machinery. Across the ToL, the number of proteins per proteome expanded roughly 200-fold from ~700 proteins in the simplest free-living DPANN (Dombrowski et al., 2019) archaea, to ~120,000 proteins in human (**Data S2**). Proteome size is similar across prokaryotes, in the order of 3000 proteins, although the smallest proteomes belong to the earliest diverging free-living DPANN archaea and *Aquificae* bacteria (Battistuzzi et al., 2004; Dombrowski et al., 2019) (not counting parasites and symbionts). Eukaryote proteomes are substantially larger, and considering free-living organisms only, the smallest eukaryotic proteomes harboring ~10,000 proteins belong to *Amoebozoa*, one of the earliest diverging eukaryotes (Burki et al., 2020; Simpson et al., 2002). Land plants and metazoans comprise hundreds of thousands of proteins per proteome. However, as described later, this dramatic increase in the number of proteins in eukaryotes occurred not only by duplication of pre-existing proteins but also by the emergence of completely new domains and folds.

#### *The median protein length increased ~2-fold* (Figure 1C)

The longer the polypeptides are the more prone they are to misfold and aggregate instead of readily reaching their native functional state (Henzler-Wildman and Kern, 2007; Mayer, 2010). Compared to ~250 residues across prokaryotes, median protein length increased about 2-fold in multicellular eukaryotes, with ~400 residues in plants and ~500 residues in Chordata (**Data S2**). Longer proteins were found primarily in multicellular eukaryotes (average lengths of the top 10% largest proteins were roughly 1,300 residues in plants, ~1,500 residues in metazoan, and ~2,150 residues in mammals). The longest polypeptides in mammals are predominantly muscle proteins, including different variants of titin (> 34,000 residues) or adhesins (> 5000 residues).

There are different ways by which proteins can increase in size. Firstly, the domains themselves can grow larger by decorating an ancestral core domain with additional segments. Secondly, the fusion of multiple domains can result in a larger multi-domain protein. Thirdly, domain-flanking regions (C- and N-termini segments, and inter-domain linkers), that are typically disordered, can expand. A systematic analysis of 38 distinct folds that are conserved across the ToL (including parasites and symbionts) showed that lengths of individual domains increased mildly, nearly 1.5-fold, across the ToL (**Figure S1A, Data S2**). The expansion of domain-flanking segments was also modest, nearly 3-fold, from prokaryotes to multicellular eukaryotes (**Figure S1B, Data S2**). Indeed, as elaborated below, length expansion primarily stemmed from the increase in the fraction of multi-domain proteins.

#### *Multi-domain proteins expanded ~50-fold* (Figure 1D)

Multi-domain proteins are inherently more prone to misfolding and aggregation than single-domain proteins, and may therefore demand more chaperone holding-unfolding-refolding action (Gong et al., 2009; Lafita et al., 2019; Liu et al., 2019). Across the ToL, multi-domain proteins comprising ≥ 3 Pfam-annotated domains have expanded ~50-fold, from ~100 proteins per proteome in prokaryotes to ~5000 in plants and animals (**Data S2**). Further, multi-domain proteins have expanded beyond the expansion of proteome size, to become nearly 3-fold more frequent in eukaryotic proteomes compared to prokaryotes (**Figure S1C**) with the expected corresponding shrinkage of proteins comprising 1□2 domains, **Figure S1D**). As described later, this expansion occurred not only by duplication of pre-existing multi-domain proteins but foremost by the emergence of new domain combinations.

### Proteome expansion by innovation

Most proteins emerge by duplication and divergence of a pre-existing protein. The outcome is paralogous proteins with the same overall fold and domain arrangement (for multi-domain proteins). Thus, duplication and ‘local’ divergence (point mutations and short insertions or deletions) certainly increase proteome size (total number of proteins) but do not dramatically change proteome composition or complexity. The latter relates primarily to the birth of completely new proteins possessing new folds, and to the emergence of multi-domain proteins with new fold-combinations. Additions of new folds and fold-combinations likely impose an additional burden on the chaperone machinery. We thus analyzed additional proteome parameters that represent expansion by innovation, rather than by mere duplication and divergence, as detailed below.

#### *Fold-types expanded ~5-fold* (Figure 2A)

To assess the emergence of new folds, we used ECOD—a hierarchical classification of protein folds that uses both sequence and structural similarities and clusters all domains with known structures into independently evolved lineages, termed as the ‘X-groups’ (Schaeffer et al., 2016). We found that from prokaryotes to eukaryotes, the number of unique folds per proteome expanded about 5-fold, from ~150 in prokaryotes to ~450 in metazoa (**Data S3**).

**Figure 2.**
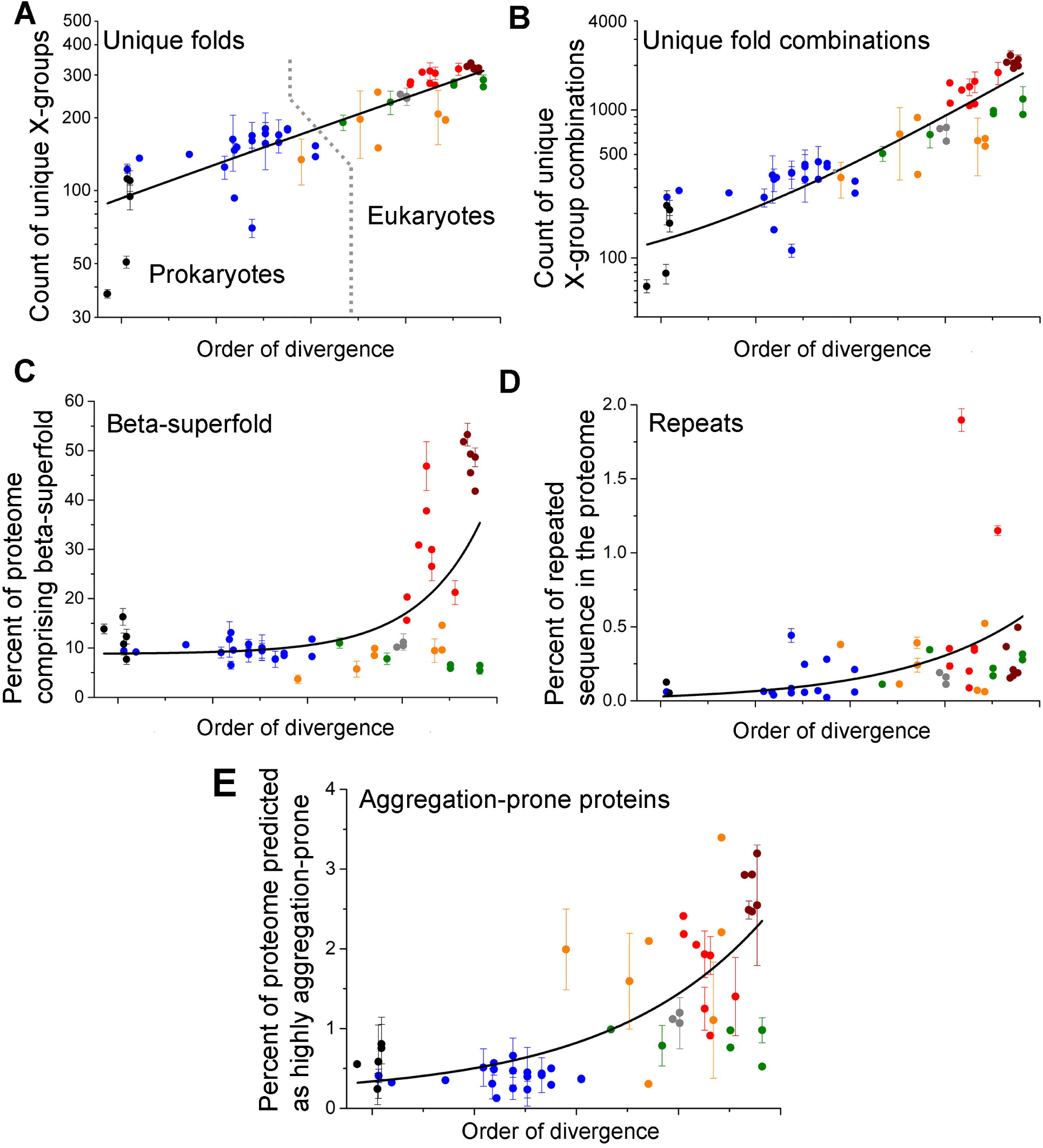
Expansion of proteomes by innovations. Figure features follow those of **Figure 1. (A)** Shown on the Y-axis (log-scale), is the average number of unique folds (Evolutionary Classification of Domains, ECOD X-groups (Schaeffer et al., 2016)) in each phylogenetic clade. The lines were derived by a fit to an exponential equation, and are provided merely as visual guides. Prokaryotic (black and blue dots) and eukaryotic organisms (orange, grey, green, red, and wine dots) were largely separated by a dashed line in panel-A. **(B)** Same, for the count of unique fold combinations. **(C)** Same, for the percent of proteins in the proteome comprising at least one beta-superfold domain (note the linear-scale). **(D)** Same, for the percent of total proteome length that is repeated (also on a linear-scale). **(E)** Same, for the percent of proteins in the proteome predicted to have ≥20 aggregation hotspots per proteome (also on a linear-scale; see also **Data S10**).

#### *New fold-combinations expanded ~20-fold* (Figure 2B)

As shown above, multi-domain proteins expanded nearly 3-fold (their fraction out to the total number of proteins) alongside a parallel shrinkage of proteins comprising 1□2 domains (**Figure S1 C, D**). This expansion of multi-domain proteins occurred not only by duplication, namely by amplifying pre-existing multi-domain proteins but also via the emergence of new combinations. To assess the latter, we examined the number of unique combinations of domains per proteome, with domains being assigned by ECOD X-groups. It appeared that new domain combinations arose throughout evolution, and from prokaryotes to eukaryotes, the number of unique combinations per proteome increased ~20-fold, from ~100 combinations in bacteria to ~2000 combinations in Chordata (**Data S4, S5**).

#### *All-beta and beta-rich folds expanded up to ~600-fold* (Figure 2C)

Proteins that are beta-rich are known to be prone to misfolding and aggregation (Pawar et al., 2005). In the simplest free-living bacteria and archaea, proteins comprising the ancient all-alpha and alpha-beta architectures are the most frequent. Remarkably, upon the emergence of eukaryotes, and in metazoans especially, all-beta or beta-rich architectures (beta-superfold) expanded massively (**Figure S2**) and became nearly 6-fold more frequent in mammalian proteomes, as compared to bacteria and archaea (*i.e*., in proportion to the total number of proteins, **Data S6**). The immunoglobulin fold had a major contribution to this expansion, owing to its diverse roles in immunity, multi-cellularity, and signaling (Halaby and Mornon, 1998).

#### *Repeat sequences expanded ~700-fold* (Figure 2D)

Proteins comprising tandem repeats of nearly identical sequences emerge readily yet are prone to misfolding and aggregation. We identified proteins with repeated sequences of the size of a single ‘foldon’ unit, ~20 aa (Englander and Mayne, 2014), with ≥ 90% sequence similarity (**Data S7**). Most of the early-diverging archaea and bacteria do not possess repeat proteins. Indeed, repeat proteins appear in more recently diverged prokaryotes and foremost in eukaryotes (a similar trend was described in (Andrade et al., 2001)). Metazoan proteomes contain large proteins with long repeated segments, *e.g. Drosophila* Ank2p (21 Ankyrin repeats, in total 836 residues), or human Dmbt1p (11 cysteine-rich repeats of total 1419 residue length). The cumulative length of repeat sequences in metazoan proteomes can be up to 100,000 residues. Overall, from prokaryotes to eukaryotes, a 700-fold expansion of repeated sequences was observed along the ToL (**Figure S3, Data S2**). Repeated sequences also expanded beyond the expansion of proteome size – the percent of total proteome length that comprises repeats increased nearly 7-fold from prokaryotes to metazoans (sum of repeat lengths normalized by the sum of all protein lengths; **Figure 2D**).

#### *Proteins predicted as aggregation-prone became ~6-fold more frequent in the proteome* (Figure 2E)

To further examine the expansion of aggregation-prone proteins, for each representative organism, we identified how many proteins in the proteome are predicted to have an unusually high number of aggregation ‘hotspots’ using the CamSol method (Sormanni et al., 2017). We then calculated what percent of the entire proteome these aggregation-prone proteins represent (see **Methods**). An ‘aggregation hotspot’ was defined as a poorly soluble protein segment of ≥5 aa length, with solubility predicted from the sequence (Sormanni et al., 2017), and the threshold for comparison was set at ≥20 hotspots per protein (at this threshold, ≤3% of proteins are aggregation-prone; the same trend was observed with lower thresholds, see **Data S10**). This prediction is restricted by the fact that some of the predicted segments actually reside in the hydrophobic cores of stably folded proteins and hence do not comprise aggregation hotspots (Sormanni et al., 2015). However, such segments are likely to be as frequent in prokaryote and eukaryote proteomes (or even less frequent in the latter where disordered proteins are abundant). Overall, the CamSol prediction indicates that compared to prokaryotes, aggregation-prone proteins have become nearly 6-fold more frequent in Chordata proteomes (**Figure 2E, Data S2**).

Overall, it appears that although gene and whole-genome duplications dominate, and in particular along with the evolution of eukaryotes (Van de Peer et al., 2009), dramatic changes in proteome composition have occurred owing to *bona fide* innovations. Specifically, concerning the burden on the chaperone machinery, proteome compositions have changed massively with respect to new folds, beta-superfolds, repeat proteins, fold combinations, and aggregation propensity.

### The evolutionary history of chaperones

In parallel to estimating the expansion of proteome size and composition, we investigated the evolutionary history of chaperones, aiming to date their emergence and their expansion along the ToL. To that end, absence or presence, and copy numbers, of the core-chaperones (HSP20, HSP60, HSP70, HSP100, and HSP90) was determined for the representative proteomes. Subsequently, protein trees were generated and compared with the ToL, to account for gene loss and horizontal transfer events. Protein sequences of all core-chaperones families were extracted from the representative proteomes (**Data S8**). These sequences were aligned and were used to generate maximum likelihood, midpoint-rooted protein trees which were then compared with the ToL (see **Methods, Table S1**).

#### *The core-chaperones emerged in early-diverging prokaryotes* (Figure 3A)

Our analysis traced the origin of all five core-chaperone families in early-diverging prokaryotes. The phylogenetic tree of a single protein, typically of a few hundred amino acids length, often lacks the resolution required to reliably date the emergence, especially when horizontal transfer events are frequent. Dating emergence to LUCA is particularly challenging. We followed the recommendations of Berkemer and McGlynn (Berkemer and McGlynn, 2020) and demanded that for a chaperone family to be assigned to LUCA, there must be a single split between bacterial and archaeal sequences at the root of the protein tree, with strong bootstrap support for this split, and that inter-kingdom branches would be longer than the intra-kingdom branches. These criteria assigned the emergence of only one core-chaperone, HSP60—a cage-like ATP-fueled unfoldase (Finka et al., 2016; Saibil, 2013), to LUCA (**Figure 3A, Table S1**). The protein tree of HSP60 further indicated an ancient horizontal gene transfer (HGT) from archaea to Firmicutes, previously noted by (Techtmann and Robb, 2010).

**Figure 3.**
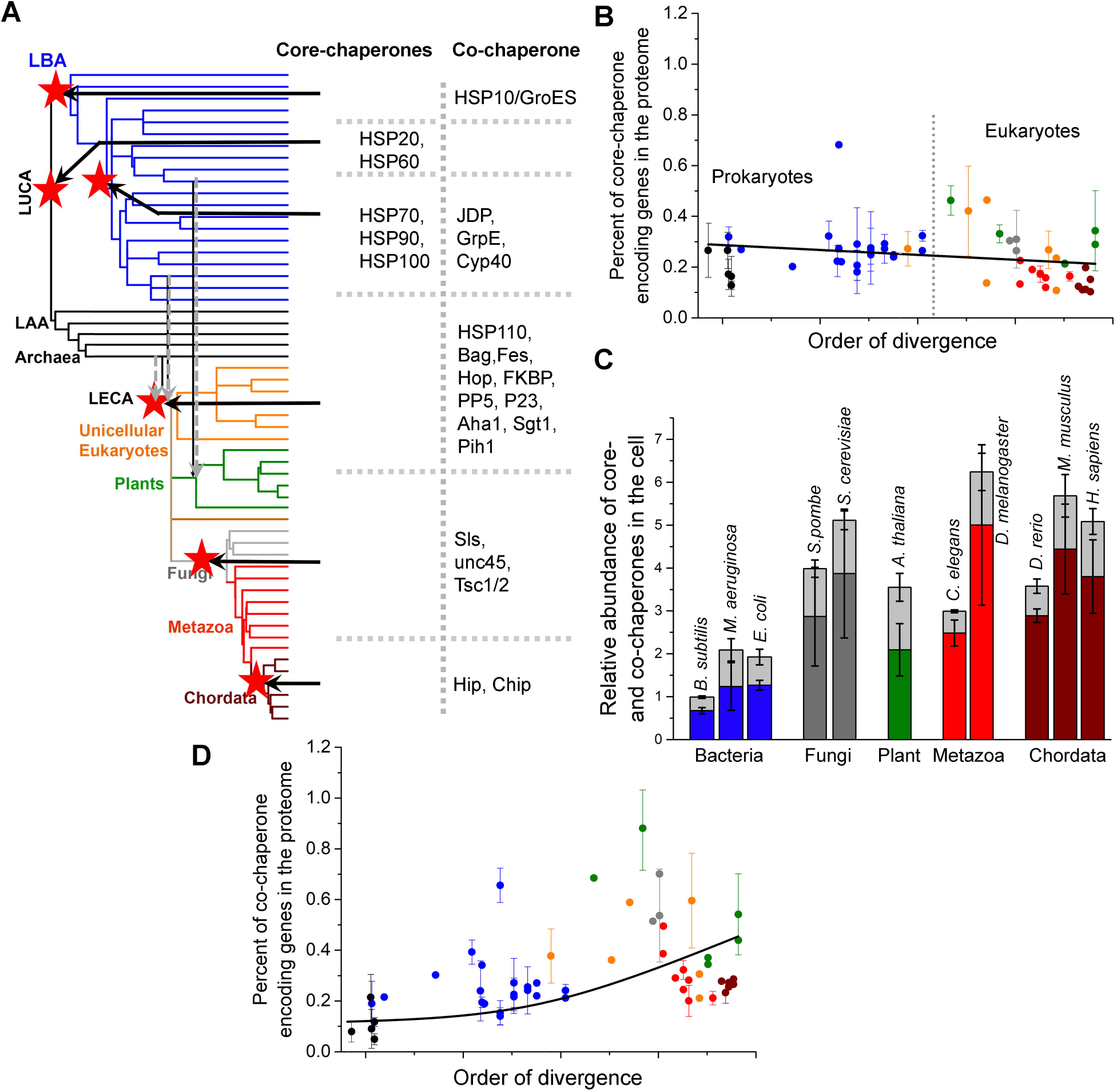
Evolutionary history of core- and co-chaperones. **(A)** The *de novo* emergence of core- and co-chaperone families are summarized on the ToL. The ToL is the same as **Figure 1A**, clade names are omitted for clarity. Ancestral nodes in which a chaperone family emerged are marked with red stars and the core- and co-chaperone families emerged in that node are listed. The dashed grey arrows reflect the endosymbiotic integration of archaeal and bacterial chaperone systems in LECA and eukaryotic and cyanobacterial chaperone systems in photosynthetic algae. **(B)** The percent of core-chaperone genes per proteome (shown is the average percentage for each phylogenetic clade). Figure features follow **Figure 1**. The line was derived by a fit to a linear equation and is provided merely as a visual guide. Prokaryotic (black and blue dots) and eukaryotic organisms (orange, grey, green, red, and wine dots) were largely separated by a dashed line. **(C)** The relative basal abundance of core- and co-chaperones in cells. Plotted are model organisms for which sufficient, reliable data were available. The columns include core-chaperones at the bottom (colors represent the phylogenetic clade in **Figure 1A**) and co-chaperones at the top (light grey color). The error bars represent the standard deviation among different non-redundant abundance datasets. **(D)** Same as panel **C** for co-chaperone genes per proteome. The line was derived by a fit to an exponential equation and is provided merely as a visual guide.

The protein tree of HSP20—an anti-aggregation ‘holding’ chaperone depicted a clear single split of bacterial and archaeal domains at the root, albeit with weak bootstrap support, and inter-kingdom branch lengths were shorter than intra-kingdom branch-lengths (**Table S1**). Similar uncertainties were noted for the majority of protein families formerly assigned to LUCA (Berkemer and McGlynn, 2020). Previous studies assigned HSP20 to LUCA (Sousa et al., 2016; Weiss et al., 2016), which, despite the above uncertainties, we concur (**Figure 3A**). Although later emergence and horizontal gene transfer (HGT) is an alternative. Indeed, in accordance with a previous study (Kriehuber et al., 2010), the HSP20 protein tree suggests multiple HGT events, though given the weak bootstrap supports it was difficult to distinguish between phylogenetic uncertainty and actual HGT events, let alone to assign donor and acceptor clades.

The remaining core-chaperone families, HSP70, HSP90, and HSP100 appear to have emerged in bacteria, though phylogenetic uncertainties and probably extensive horizontal transfer between different bacterial clades, and between bacteria and archaea, prevent the reliable assignment of their points of origin (**Figure 3A**). The ATP-dependent core-chaperone HSP70—that controls protein unfolding, disaggregation, translocation, and degradation (Fernandez-Fernandez and Valpuesta, 2018; Rosenzweig et al., 2019), was detected in the earliest diverging bacterial clades *Aquificae* and *Thermotogae*. However, the protein tree clustered their HSP70 sequences with those from later-diverging bacterial lineages *Deltaproteobacteria, Clostridia*, and *Bacilli*, with bootstrap values being too low to distinguish between phylogenetic uncertainty and true HGT events (**Table S1**). Thus, HSP70 appears to have a bacterial origin, around or after the emergence of terrestrial bacteria, *i.e*., around the divergence of *Fusobacteria* (Battistuzzi and Hedges, 2009) (**Figure 3A**). Following its emergence, HSP70 was likely horizontally transferred to archaea, but the current analysis could not reliably assign donor and acceptor clades.

The protein trees of both HSP90 and HSP100 depict a similar scenario. Both seem to have emerged in bacteria around or after terrestrial bacteria emerged (**Figure 3A, Table S1**). Although a reliable point of origin could not be assigned for either of these two chaperones, biochemical assays show that whereas the activity of HSP90 or HSP100 strictly depends on the presence of HSP70, HSP70 itself can act independently. Further, across the ToL, every organism that harbors genes for HSP90 and or HSP100 also harbors genes for HSP70, but not *vice versa*. Thus, it is likely that HSP90 and HSP100 have both emerged after HSP70. Similar to HSP70, HSP90 and HSP100 were likely horizontally transferred to archaea. While our protein trees do indicate such trends, the bootstrap values are low.

The archaeal and bacterial core chaperones were integrated into LECA, and no new core-chaperones emerged with the birth of eukaryotes (**Figure 3A**). Chaperones of archaeal origin mostly continued to function in their original compartment, the cytosol. Although most *Alphaproteobacterial* endosymbiont genes were transferred to the nucleus, most of the chaperones of bacterial origin evolved to translocate back to the compartment from which they originated, namely to the mitochondria (Bogumil et al., 2014; Hemmingsen et al., 1988; Kampinga et al., 2019; Martin et al., 2002). Chaperone evolution in eukaryotes involved gene loss as well, *e.g*., cytosolic and mitochondrial HSP100 have been lost in metazoa (Kampinga et al., 2019).

#### *The expansion of core-chaperones* (Figure 3B)

Whereas no new core-chaperone family emerged in eukaryotes, gene copy numbers of the existing families did increase via gene duplication, to support expanding proteomes, for condition-specific expression, and also to cater for the emergence of multiple subcellular compartments. Accordingly, bacteria and archaea that possess the same five core-chaperone families as eukaryotes, copy numbers of individual families per proteome range between 1–4, summing up to an average of ~8 core-chaperone genes per proteome (**Figure S4, Data S8**). In comparison, the number of core-chaperone genes in higher plants, which are among the most complex eukaryotes, increased ~30-fold for HSP20 (**Figure S4A**), ~50-fold for HSP60 (**Figure S4B**), ~40-fold for HSP70 (**Figure S4C**), ~20-fold for HSP90 (**Figure S4D**), and ~10-fold for HSP100 (**Figure S4E**). Parasitic microbes, such as *Mycoplasma pneumoniae, Plasmodium falciparum*, and *Entamoeba histolytica*, and photosynthetic bacteria, algae, and plants often harbor unusually high chaperone gene copy numbers, likely to counter the immune response of the host (Day et al., 2019; Shonhai et al., 2011) and the oxidative stress (Dahl et al., 2015; Niforou et al., 2014). However, when proteome size is accounted for, it is evident that the expansion of core-chaperones largely coincides with the overall expansion of proteome size. In fact, core-chaperones comprise ~0.3%, *i.e*., 3 out of 1000 proteins, in all proteomes, from the simplest free-living prokaryotes to mammals (**Figure 3B**). Further, the expansion of core-chaperones occurred by gene duplication only, with no *bona fide* innovation, as all five core-chaperone families seem to have preexisted in prokaryotes.

#### *Core-chaperones became ~4-fold more abundant in the cell* (Figure 3C)

As described above, the frequency of core-chaperone genes remained roughly constant in both prokaryotes and eukaryotes. However, gene expression levels could vary, and increased protein abundance could support the increasingly complex proteomes of eukaryotes. To examine this, the relative basal abundance of core- or co-chaperones in the cell was compared. Unfortunately, this analysis is restricted to model organisms in which ample proteomics data are available. Non-redundant proteome-scale abundance data was collected for 11 model organisms spanning 9 major clades along the ToL (see **Methods, Data S9**). As plotted in **Figure 3C**, the relative basal abundance of core-chaperones in the cell, compared to all other proteins, appears to have increased ~4 fold across the ToL, from an average of 1.1% in bacteria (with ~0.7% in Firmicutes, the earliest diverging prokaryote clade for which reliable proteomics data are available) to an average of 4.2% in Chordata.

#### *Co-chaperones expanded ~9-fold* (Figure 3D)

In eukaryotes, as the gene copy numbers of core-chaperones expanded, and their protein abundance also increased, what happened to their auxiliary workforce, the co-chaperones? The number of unique co-chaperone families per proteome expanded from ~3 in prokaryotes to ~20 in humans. Most co-chaperones are eukaryote-specific (**Data S8**), and therefore have likely emerged relatively recently, and only a few co-chaperones are found in all three domains of life. To date their emergence, protein trees were generated. These suggest that, as expected, these co-chaperones emerged after the core-chaperone they work with. HSP60 is assigned to LUCA, while its bacterial co-chaperone HSP10/GroES appears to have emerged later along with the emergence of bacteria (**Figure 3A, Table S1**). HSP70’s co-chaperones, the J-domain proteins (JDPs) and GrpE, and HSP90’s co-chaperone Cyp40, appear to have emerged at the same node as their respective core-chaperones. Given the phylogenetic uncertainties, possible extensive horizontal transfers, and the resolution that protein trees allow, a more precise dating could not be performed. It appears, however, that JDP, GrpE, and Cyp40, have all emerged after the emergence of terrestrial bacteria (**Figure 3A**), and therefore, likely emerged after their respective core-chaperones. Several co-chaperone families emerged in eukaryotes (**Figure 3A**), including HSP110 that diverged by duplication of HSP70 (Sarkar et al., 2013), and Pih1, Aha1, and Chip that harbor eukaryote-specific folds and hence likely emerged *de novo*. Overall, it is evident that co-chaperones tail core-chaperones, and not *vice versa*. With the birth of several co-chaperones in eukaryotes, the percentage of genes encoding for co-chaperones in the proteome expanded ~9-fold from prokaryotes to eukaryotes (**Figure 3D**). Notably, the JDPs, co-chaperones of HSP70, are the major contributor to this copy-number expansion (Bogumil and Dagan, 2012).

Our analysis, therefore, suggests that evolutionary innovation occurred primarily at the level of co-chaperones that facilitated the basal core-chaperone activity thus expanding the chaperone network to meet the challenges of newly emerging protein folds and increasingly complex proteomes.

## Discussion

How did chaperones evolve to support proteome complexity? To address this question, we compiled data from multiple sources and analyzed them under one roof, thus allowing a systematic, quantitative comparison, as summarized in **Figure 4**. From the simplest free-living prokaryotes to plants and animals, proteomes have continuously expanded by both duplications and innovations. It is primarily due to the latter that proteome ‘complexity’ has continuously increased in various ways that demand increased chaperone action. Across the ToL and especially when comparing prokaryotes to eukaryotes, we see a larger number of proteins per proteome (**Figure 4A**, light grey bars) as well as larger proteins (light yellow bars). The latter relates to multi-domain proteins being increasingly represented (grey bars). The number of different folds increases as well as of unique combinations of folds in multi-domain proteins (yellow and orange bars, respectively). Proteomes also contain a larger fraction of protein types that are prone to misfolding, such as repeat proteins (dark grey bars) and proteins comprising beta-sheets (wine bars), and those that are predicted to be highly aggregation-prone (dark yellow bars).

**Figure 4.**
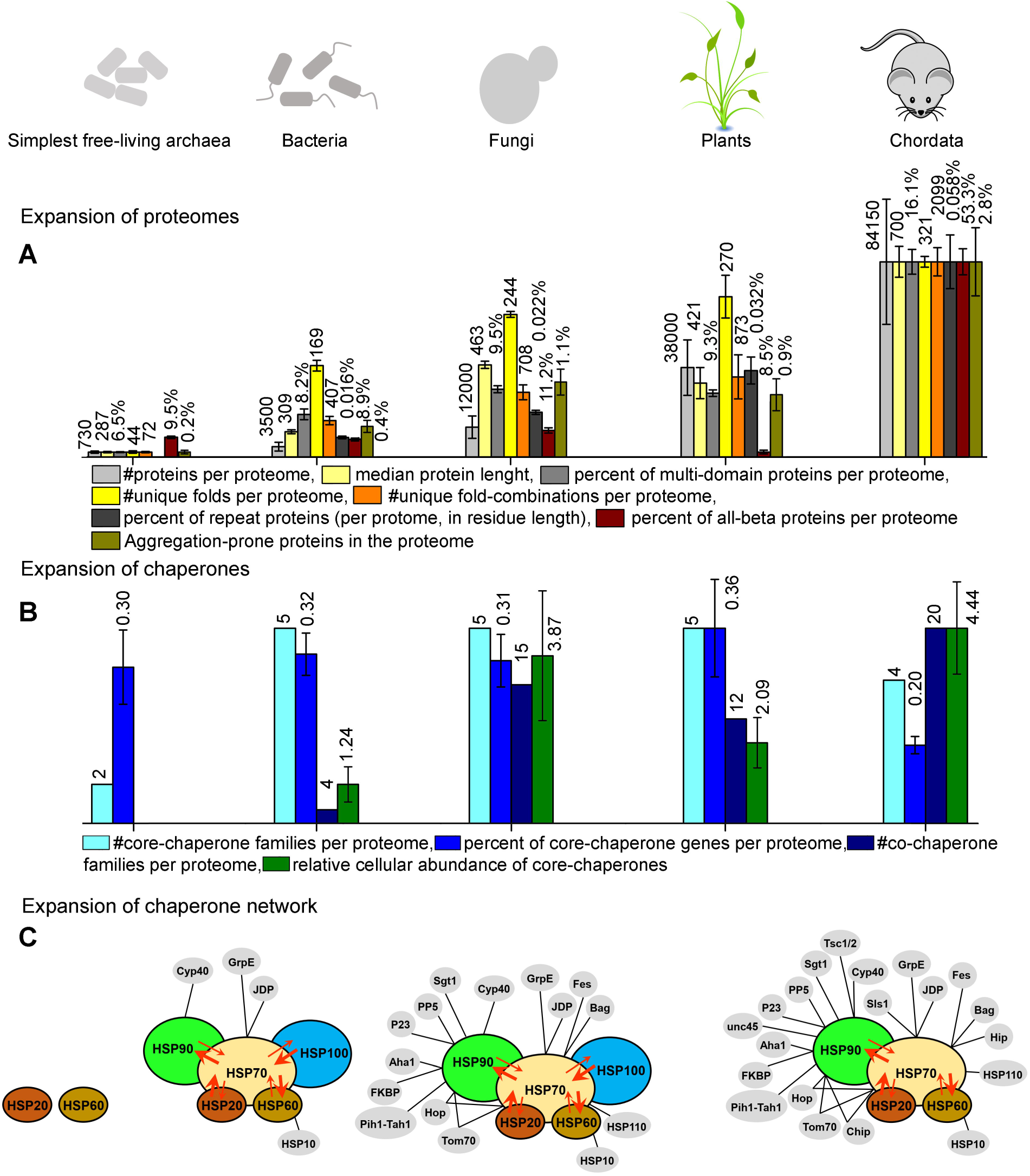
A summary figure describing the parallel expansion of proteomes and chaperones. Bar heights (Y-axis) were scaled such that the highest value per each parameter assumed the same height (the absolute values are listed above the bars). **(A)** Bar graphs describing the expansion of proteomes in a nutshell. For the simplest free-living archaea, bacteria, fungi, plants, and chordates, plotted are the number of proteins per proteome (light gray), the median protein length (light yellow), the number of unique folds (grey), the number of unique fold-combinations per proteome (yellow), the percent of multi-domain proteins (out of all proteins in the proteome; orange), the percent of proteome length that corresponds to repeat proteins (calculated by residue length; dark grey), the percent of proteins that have the beta-superfold architecture (wine), and the percent of proteins predicted as highly aggregation-prone (dark yellow).. **(B)** Same, for the expansion of chaperones. Plotted are the number of core- (cyan) and co-chaperone families per proteome (navy), the percent of core-chaperone genes in the proteome (blue), and the relative cellular abundance of core-chaperones compared to all other proteins (green). **(C)** A schematic description of the expansion of the integrated chaperone network. Core chaperones are shown in various colors and with black outlines, while co-chaperones are in grey with no outline. Co-chaperones of HSP60, HSP70, and HSP90 are connected to their respective core-chaperone by black lines. Cooperativity between core-chaperones is represented by overlaps between circles, and substrate sharing between different core-chaperones are shown by red arrows. Arrow direction and width represent the direction and magnitude of substrate sharing. Note that the network is shown for the simplest free-living archaea, bacteria, fungi, and chordates.

This dramatic increase in proteome complexity, and hence in the demand for chaperone action, has not been met by the emergence new of core-chaperones. Eukaryotes possess the same five core-chaperone families as prokaryotes and metazoans and chordates have in fact lost HSP100 (**Figure 4B**, cyan bars). Further, the relative representation of core-chaperone genes does not vary between prokaryotes and eukaryotes (blue bars). Rather, the need for increased chaperone action was met in two ways. Firstly, by an increased cellular abundance of core-chaperones (green bars). Secondly, although, by the emergence of new co-chaperones—while in bacteria ~4 co-chaperone families are found, in eukaryotes their number increased to 15 or even 20 in Mammalia (navy blue bars).

The above trends highlight two features that comprise hallmarks of the chaperone machinery. The generalist nature of core-chaperones, and their ability to act in a cooperative mode alongside co-chaperones as an integrated network. HSP60, HSP70, HSP90, and HSP100 are core-chaperones acting as generalist unfolding-refolding machineries that work on a broad range of differently misfolded and aggregated protein substrates, largely regardless of size (except HSP60 (Bukau and Horwich, 1998)), structure and function (Finka et al., 2016; Gong et al., 2009). While core-chaperones can exert high-affinity binding to few specific substrates at their native folded state, they generally tend to bind misfolded and aggregated polypeptides that abnormally expose hydrophobic surfaces (Gong et al., 2009; Rudiger et al., 1997; Scheibel et al., 1998). The main driving force for duplication and specialization is a functional tradeoff—optimization of one function comes at the expense of other functions (Tawfik and Gruic-Sovulj, 2020). However, given a ‘generalist’ mode of function, the quality-control of increasingly large and complex proteomes could be achieved by an increase in abundance of core-chaperones, rather than by the emergence of new core-chaperone families. Indeed, although gene copy-numbers of core-chaperones have indeed increased by gene duplication, their relative representation compared to proteome size remained constant (**Figure 3B**) and the resulting paralogous copies have mostly re-localized to different subcellular compartments or are expressed under different stress conditions (Kampinga et al., 2009). In parasitic microbes and photosynthetic organisms, duplicates of HSP70 and HSP90 have sub-specialized to resist host immune responses and oxidative stress (Dahl et al., 2015; Day et al., 2019; Niforou et al., 2014; Shonhai et al., 2011). However, consistent with their generalist nature, the challenge of maintaining large, complex proteomes (**Figure 4A**) has primarily been met by an increased abundance of preexisting core-chaperones rather than by the *de novo* emergence of new ones. This is also reflected in their generic name ‘heat-shock proteins’ – under stress conditions, cells use the very same core-chaperones, yet just elevate their abundance.

In healthy cells, an integrated chaperone network, comprising both core- and co-chaperones, controls protein quality (Hipp et al., 2019; Jayaraj et al., 2020; Taipale et al., 2014). In this network (**Figure 4C**), the highly abundant core-chaperones operate cooperatively, namely, they not only share, and exchange incompletely processed misfolded or unfolded protein substrates, but also trigger the activities of one another. HSP70 plays a critical role in this network by mediating cooperative communications between the other core-chaperones. For example, HSP70 triggers the disaggregase activity of HSP100, and jointly they disaggregate aggregated proteins and promote their subsequent refolding (Glover and Lindquist, 1998; Haslberger et al., 2008; Rosenzweig et al., 2013). In another example, HSP20 can transfer misfolded substrates to HSP70 for ATP-driven unfolding, from which they can be further transferred to HSP60 for final refolding to the native state (Veinger et al., 1998). Likewise, HSP90 can promote the maturation of incompletely processed HSP70-substrates (Genest et al., 2011; Moran Luengo et al., 2018). Cooperativity and substrate-sharing between core-chaperones are schematically represented in **Figure 4C**. Together, these generalist, cooperative core-chaperones constitute the core of an integrated chaperone network that has emerged from a simple two-component system in LUCA (**Figure 4C**).

Alongside the expansion of proteome complexity, the chaperone network has also expanded— primarily by the emergence of co-chaperones (**Figure 4C**). This expanding array of co-chaperones augmented the ability of core-chaperones to efficiently share substrates and to function cooperatively. In contrast to the generalist core-chaperones, co-chaperones are more diverse and accordingly seem to sub-specialize in specific roles, including co-chaperones that handle specific proteins. Examples include UNC45, a co-chaperone that emerged in Fungi, and facilitates HSP90-mediated maintenance of myosin in metazoan skeletal and cardiac muscles (Wohlgemuth et al., 2007). Another Fungi-born co-chaperone, Tsc1/2 heteromer, specializes in recruiting kinase and some non-kinase substrates to HSP90 (Woodford et al., 2017). Other co-chaperones mediate protein transport; examples include Tom70 and P23 that facilitate protein trafficking through Golgi and mitochondrial membranes (Craig and Marszalek, 2017; Echtenkamp et al., 2011; Melnyk et al., 2015). The specialist mode of function of co-chaperones coincides with how they expanded, namely by duplication and divergence of ancient prokaryote-born co-chaperones, but also via *bona fide* innovations, *i.e*., by the emergence of completely new specialized co-chaperones in eukaryotes. As shown here, the emergence of new co-chaperones coincides with the emergence of new proteins (*i.e*., by de novo emergence rather than by duplication of pre-existing proteins). However, co-occurrence does not mean co-evolution—indeed, we know very little about the latter. Did certain co-chaperones emerge to support the de novo emergence of a specific protein or protein class? If so, does chaperone dependency persist, hence making co-dependency a ‘selfish’ irreversible trait? Alternatively, as some newly-emerged proteins evolved further, their foldability improved, allowing them to become chaperone-independent.

Thus, across the tree of life, proteomes have massively expanded, not just by duplication of pre-existing proteins but also by the emergence of completely new ones. Eukaryotic proteomes became particularly large and specifically richer in repeat, beta-rich, and aggregation-prone proteins whose folding is inherently challenging. These changes in proteome size and composition intensified the demand for chaperone action. Curiously, however, no new core-chaperones emerged in response to this increased demand. Instead, they increased in abundance relative to all other proteins in the cell. Foremost, an entire network of co-chaperones had evolved that facilitate the basal core-chaperone activity.

## Supporting information

Response to Reviewers

## Author Contributions

S.M., M.R., P.G., and D.S.T. conceived and conceptualized the research. M.R. performed the phylogenetic analysis shown in Figure 3A. M.R. and S.M. performed the chaperone copy number analysis shown in Figure 3B, 3D, and 4B. S.M. performed the Tree of Life and proteome expansion analysis shown in Figures 1, 2, and 4A, and the chaperone abundance analysis shown in Figures 3D and 4B. S.M. also prepared all the Figures and wrote the original draft. S.M., P.G., and D.S.T. reviewed and edited the manuscript. D.S.T. and P.G. acquired funding and supervised the study.

## Acknowledgment

We sincerely thank Jagoda Jablonska for helping to establish the representative ToL, and to Ita Gruic-Sovulj for proposing the chaperone abundance analysis. We thank Rina Rosenzweig, Harm H. Kampinga, Matthias P. Mayer, Sudip Kundu, and Agnieszka Klosowska for valuable suggestions regarding the manuscript.

## Supplementary Figure Legends

**Figure S1** (related to Figure 1). Figure features follow those of **Figure 1. (A)** Shown on the Y-axis (linear-scale), are the average lengths of the protein domains that belong to 38 ECOD X-groups present throughout the ToL.

**(B)** Same, for the length of the domain-flanking segments.

**(C)** Same, for the percent of proteins in the proteome comprising ≥ 3 Pfam-annotated domains.

**(D)** Same, for the percent of proteins in the proteome comprising < 3 Pfam-annotated domains.

**Figure S2** (related to Figure 2). Figure features follow those of **Figure 1. (A)** Shown on the Y-axis (log-scale), is the number of beta-superfold proteins in the proteome.

**Figure S3** (related to Figure 2). Figure features follow those of **Figure 1. (A)** Shown on the Y-axis (log-scale), is the total length (amino acids) of repeated sequence in the proteome.

**Figure S4** (related to Figure 3). Figure features follow those of **Figure 1**. Shown on the Y-axis is the average number, per genome, of genes encoding **(A)** HSP20 (note the log-scale), **(B)** HSP60, **(C)** HSP70, **(D)** HSP90, and **(E)** HSP100.

## Supplementary Table Legends

**Table S1** (related to Figure 3). Protein trees (newick format) of core and co-chaperone families. These trees were inferred by using the Maximum Likelihood method. Initial trees for the heuristic search were obtained by NJ/BioNJ algorithm, using JTT distance model.

## Supplementary Data File Legends

**Data S1**. The 188 representative organisms and their phylogenetic classifications (related to Figure 1).

**Data S2**. Numerical values of the proteome parameters (related to Figure 1 and Figure 2).

**Data S3**. A list of all the unique ECOD X-groups, and Pfam clans, detected in representative proteomes of our Tree of Life, and a list of ECOD X-groups conserved across the ToL (related to Figure 2).

**Data S4**. A list of all the unique ECOD X-group combinations detected in the representative proteomes of our Tree of Life (related to Figure 2).

**Data S5**. Unique Pfam domain combinations, and clan combinations, detected in the representative proteomes; ‘no clan’ relates to Pfam families that are not assigned to a clan, and thus reflect an independent lineage (related to Figure 2).

**Data S6**. Expansion of beta-superfolds beyond the expansion of proteome size: for representative organisms of our Tree of Life, shown are the percent of proteins in the proteomes that comprise a domain related to one of the ECOD top hierarchies (related to Figure 2).

**Data S7**. Repeat segments identified in the representative proteomes (related to Figure 2).

**Data S8**. Pfam-annotated domain combinations and the UniProt identifiers of the core and co-chaperones identified in the representative proteomes (related to Figure 3).

**Data S9**. Relative abundance of core- and co-chaperones in 11 model organisms (related to Figure 3).

**Data S10**. The percent of aggregation-prone proteins per proteome (related to Figure 2).

## Methods

### Constructing the Tree of Life

To construct a representative Tree of life, we used the TimeTree database (Kumar et al., 2017) and the NCBI taxonomy database (Federhen, 2012). For all major clades of bacterial, archaeal, and eukaryotic domains, non-redundant representative species were selected, ensuring that (i) the minimum splitting time for any pair of taxa is ≥ 65 million years, (ii) their annotated proteome sequences are available in NCBI genome database (Benson et al., 2013) and (iii) proteome-scale domain assignment data are available in Pfam (Finn et al., 2014). This analysis rendered 188 representative organisms covering 56 major bacterial, archaeal, and eukaryotic clades (**Data S1**). Note that phylogenetic analyses often assign parasitic and symbiotic organisms that have experienced reductive evolution as the earliest diverging clades of their corresponding kingdoms of life. Examples include *Nanoarchaeum equitans*, an obligate symbiont, assigned as the earliest diverging archaea (Hedges et al., 2015; Huber et al., 2002; Waters et al., 2003), and parasitic Excavates assigned as one of the earliest diverging eukaryotes (Burki et al., 2020; Simpson et al., 2002). While these organisms were included in our representative set of organisms, the expansion of proteomes and chaperones has been analyzed for free-living organisms only.

The TimeTree Database comprises phylogenetic relationships, as well as literature-based annotations of predicted emergence times (Million Years Ago, MYA, from now) for > 50,000 species. Using this database, a Tree of Life (ToL) was constructed in which leaves represent the extant clades (*e.g*., Mammalia, which comprises three representative organisms: human, cat, and mouse). The tree’s root comprises the Last Universal Common Ancestor (LUCA), and the nodes represent the hypothetical ancestors. Branch lengths represent the divergence time of the different clades. One representative species from each of our 56 phylogenetic clades was chosen, and this species set was submitted to the TimeTree Database to extract a ‘core-tree’ for their original tree. The obtained tree topology was manually adjusted to depict the emergence of eukaryotes from Asgard archaea and *Alphaproteobacteria* by an endosymbiosis event. The branch lengths represent the evolutionary divergence times as documented in TimeTree, and were used as proxies of the order of divergence to plot the proteome parameters.

### Capturing proteome expansion

#### Proteome size and median protein length in the proteome

Annotated proteome sequences of the 188 representative organisms were obtained as FASTA-format files from the NCBI genome database (Benson et al., 2013). For each organism, we used Biopython v1.75 package to compute the total number of proteins in the respective proteome, and the length of each protein. These protein lengths were used to estimate the median protein length for each organism (**Data S2**).

#### Multi-domain proteins in the proteome

Proteome-scale domain annotations of each of the 188 representative organisms were obtained from Pfam (Finn et al., 2014). In Pfam, the established profile HMMs of known domain families (Hidden Markov Models, probabilistic models used for the statistical inference of homology) are searched against the protein sequences, to find all instances of that domain. A statistical significance score (the probability that the prediction is a random hit) is assigned to each predicted instance based on the sequence similarity with the profile HMM. Any domain assigned with *p* < 10^−5^ significance were considered for further analysis. Overall, in these 188 representative organisms, we identified 7694 domain families, which amounts to roughly 43% of all annotated domain families (17836 families) in Pfam 32.0, September 2018 release. For each species, the number of proteins comprising <3 and those comprising ≥3 Pfam-annotated domains were counted (**Data S2**).

#### Number of unique fold types and fold-combinations

In general, sequence homologies between different domain families show that they can be clustered into independently evolved lineages (meaning, members of different lineages do not exhibit any detectable sequence homology). Two databases, Pfam (Finn et al., 2006) and ECOD (Schaeffer et al., 2016) perform this clustering considering sequence and structural similarities among domain families as the benchmark. In Pfam, these independently evolved super-families are remarked as Clans (Finn et al., 2006). In ECOD, this clustering is hierarchical (from bottom to top: F-group, T-group, X-group, and top hierarchy) and the independently evolved lineages are termed as the X-groups (Schaeffer et al., 2016). For each of the 188 proteomes, we mapped the Pfam-assigned domains to the Pfam-Clans as well as to the ECOD X-groups. The 7694 domain families were mapped to 554 unique Clans, and to 976 X-groups, covering 88% of all annotated Clans (629 Clans in Pfam 32.0), and 42% of all annotated ECOD X-groups (2316 X-groups, ECOD v20191115). The numbers of unique Clans, or X-groups, identified in a given proteome were considered as a measure of the number of unique fold types present in a given organism (**Data S3**).

This analysis further allowed us to estimate the vectorial combinations (AB ≠ BA) of Clans / X-groups, from N- to C-terminal, present in a given protein. The total number of Clan / X-group combinations present in the proteome represents the total number of unique fold-combinations present in the respective organism. The 554 Clans and 976 X-groups included in our work yielded to 15463 X-group combinations and 16538 Clan combinations respectively (**Data S4, S5**).

#### Proportion of beta-superfolds in the proteome

To capture the proportion of beta-structures in the proteome, for each representative organism, we identified how many proteins in the proteome are annotated as belonging to all-beta folds, and what fraction of the entire proteome they represent. All beta structures assigned under ECOD top hierarchies beta-barrel, beta meander, beta-sandwich, beta duplicates or obligate multimers, and beta complex topology were considered in the analysis (**Data S6**).

#### Repeated sequences in the proteome

To capture the abundance of repeated sequences in the proteome, the 188 representative proteomes were scanned by T-REKS repeat-identifier program (Jorda and Kajava, 2009). All repeats that are larger than a ‘foldon’ unit (~20 aa) (Englander and Mayne, 2014), and exhibit ≥ 90% sequence similarity were considered for further analysis (**Data S7**). Summing the lengths of all the identified repeats, we derived the total length of repeated sequences in each proteome, and what fraction of the total proteome length it covers (total repeat length normalized by the sum of all protein lengths, **Data S2**).

#### Expansion of domain lengths

To examine the expansion of domain lengths, we first identified 38 distinct folds (X-groups) that are conserved across the ToL, including parasites and symbionts (**Data S3**). Proteins harboring these X-groups were detected in the representative organisms and the respective domain lengths, as annotated in Pfam, were estimated. For each organism, these lengths were pulled together and the average domain length was estimated (**Data S2**).

#### Expansion of domain-flanking regions

For each protein harboring an X-group conserved across the ToL, we used the Pfam-annotated location of the protein domain in the primary sequence to estimate the lengths of C- and N-termini segments, and inter-domain linkers. For each organism, the lengths of these domain-flanking regions were pulled together to estimate the average (**Data S2**).

#### Predicted proportion of aggregation-prone proteins in the proteome

To capture the proportion of aggregation-prone proteins in the proteome, for each representative organism, we identified how many proteins in the proteome comprise ≥ *n* ‘aggregation hotspots’, where *n* is an integer. An ‘aggregation hotspot’ was defined as a ‘poorly soluble’ protein segment of ≥5 aa length, with solubility predicted from the protein’s sequence using CamSol v2.1 (Sormanni et al., 2017). The CamSol method yields the solubility profile of a protein: a solubility score is assigned to each amino acid in a way that regions with scores >1 denote ‘highly soluble’ regions, while scores <−1 reflect ‘poorly soluble’ ones. For each of the 188 representative organisms, we estimated the percent of proteins in the proteome that comprises ≥ *n* ‘aggregation hotspots’, for 2 ≤ *n* ≤ 20 (**Data S10**). Note that the CamSol method does not take structural information into account, which means at least for globular proteins, the predicted poorly soluble regions might reside in the hydrophobic core of globular proteins (Sormanni et al., 2017).

### Capturing chaperones in the proteome

To determine the evolutionary appearance and expansion of the core-chaperone (HSP20, HSP60, HSP70, HSP90, and HSP100), and co-chaperone families (HSP10/GroES, JDP/HSP40, HSP110, GrpE, Bag, Fes, Hip, Hop, Chip, Tom70, Cyp40, FKBP52/51, PP5, Unc45, Cdc37, P23, Aha1, Sgt1, Pih1, Tah1, and Tsc1/2 heterodimer) we identified their occurrences in the 188 representative organisms, using two complementary methods.

#### Identifying chaperones by BLAST-search

Chaperone family proteins were manually curated from the UniProt (UniProt, 2015) annotated proteomes of model organisms (*Escherichia coli* and *Saccharomyces cerevisiae*). These protein sequences were then used as queries to find orthologous sequences in the other organisms by a comprehensive protein-protein BLAST (Altschul et al., 1990). BLAST hits associated with 50% sequence coverage and ≤ 10^−5^ e-values (Marcet-Houben and Gabaldon, 2015) were manually inspected to extract the ‘true-positive’ chaperone family members.

#### Identifying chaperones by characteristic domain combinations

The second method involved manually looking into the Pfam-assigned domain combinations of annotated chaperones. For instance, bacterial HSP20s (*e.g*., IbpA protein in *E. coli*) predominantly comprise a single HSP20 domain, whereas chordate HSP20s (*e.g*., HSPB6 protein in *H. sapiens*) often comprise an additional N-terminal alpha-crystallin domain. For each chaperone family, we investigated their domain organization in Pfam and constructed a library of domain combinations (occurrence of the domain of interest with other domains in annotated proteomes). This library comprised any combination that Pfam reports in at least 10 different sequences. We searched these combinations in the 188 representative proteomes. Any protein comprising any of these domain combinations was assigned to be a member of that core-chaperone family. This analysis excluded eukaryote specific HSP110 co-chaperone family that is composed of two HSP70 domains, and thus cannot be distinguished from HSP70s that comprise two HSP70 domains. In this case, specific hallmark sequences in the linker and the more variable C-terminal domain of HSP110 that differ from HSP70 were used in BLAST searches.

The two complementary methods rendered an identical number of chaperone gene copy-numbers in each organism, reflecting the robustness of the overall approach (**Data S8**).

### Phylogenetic analysis

Multiple sequence alignments (MSAs) were generated using Muscle v3.8.31 (Edgar, 2004). The alignments were manually curated and gap-majority columns were trimmed using trimAl v1.330 (Capella-Gutierrez et al., 2009). Only for HSP20, alignments were collected from EggNOG database (Huerta-Cepas et al., 2019), and were manually curated. These alignments were filtered by removing all sequences that are more than 80% identical (usually redundant sequences) and those that are less than 30% identical (usually interferes with rooting by creating extremely long branches) were removed using T-coffee (Di Tommaso et al., 2011). Maximum likelihood phylogenetic trees were generated by MEGA X (Kumar et al., 2018), using JTT distance matrix and NJ/BioNJ initial tree. Phylogenetics trees are provided in **Table S1**. To date the emergence of individual chaperone families, the protein trees were manually compared with the ToL to assign the node of emergence and possible HGT events.

### Chaperone abundance analysis

The abundance of a protein simply means the number of copies of a protein molecule in a cell. Non-redundant genome-scale abundance data was collected for 11 non-extremophilic model organisms from PAXdb (Wang et al., 2015), measured under normal conditions. This database comprises whole genome protein abundance information across organisms and tissues. Dataset quality was measured by its proteome coverage and interaction consistency score (ICS). Proteome coverage represents what fraction of the entire proteome is included in the abundance dataset. ICS is based on the assumption that proteins that contribute jointly to a shared function (such as members of a protein complex) should tend to have roughly similar protein abundance levels. For unicellular organisms, datasets that cover ≥40% of the respective proteomes with ≥3.0 ICS and include ≥30% of all core- and co-chaperones identified in the respective organism, were selected for analysis. For multicellular organisms, tissue-specific data was collected. Datasets that cover ≥10% of the respective proteomes with ≥3.0 ICS and includes ≥30% of all core- and co-chaperones identified in the respective organism, were selected for analysis. All datasets related to stress or disease-conditions were removed and the remaining 121 datasets were considered for further analysis (**Data S9**). For each abundance dataset, we classified the list of proteins into three non-overlapping groups: (i) core-chaperones, (ii) co-chaperones, and (iii) proteins that are not chaperones. For each group, the abundance values of all the proteins were summed to estimate the ‘total abundance’. The ‘total abundance of core-chaperones’, normalized by the ‘total abundance of proteins that are not chaperones’ represents the relative abundance of core-chaperones. Similarly, we estimated the ‘relative abundance of co-chaperones’. For each species, the average and standard deviations among different datasets were measured.

### Visualizing the expansion of proteomes and chaperones

For each clade in our core-tree, numerical values of the proteome parameters and chaperone-copy number values obtained for its representative organisms were pulled together to estimate the clade average, and the clade standard deviation. The clade average values were subsequently plotted against their predicted divergence time and were fitted into the standard exponential growth curve. The fitted exponential curves are depicted as solid black lines.

### Graphics

The Tree of Life was generated using the Interactive Tree Of Life (iTOL) v4.0 (Letunic and Bork, 2019). All the plots were generated using OriginPro software v9.1.0. Figures were compiled using Adobe Photoshop CS6 v13.0.1.

### Quantification and statistical analysis

All the statistical analyses mentioned in the main text were performed using in-house Python and R scripts.

