## Supplementary figures and images for "On the evolution of chaperones and co-chaperones and the expansion of proteomes across the Tree of Life"

### Figure S1

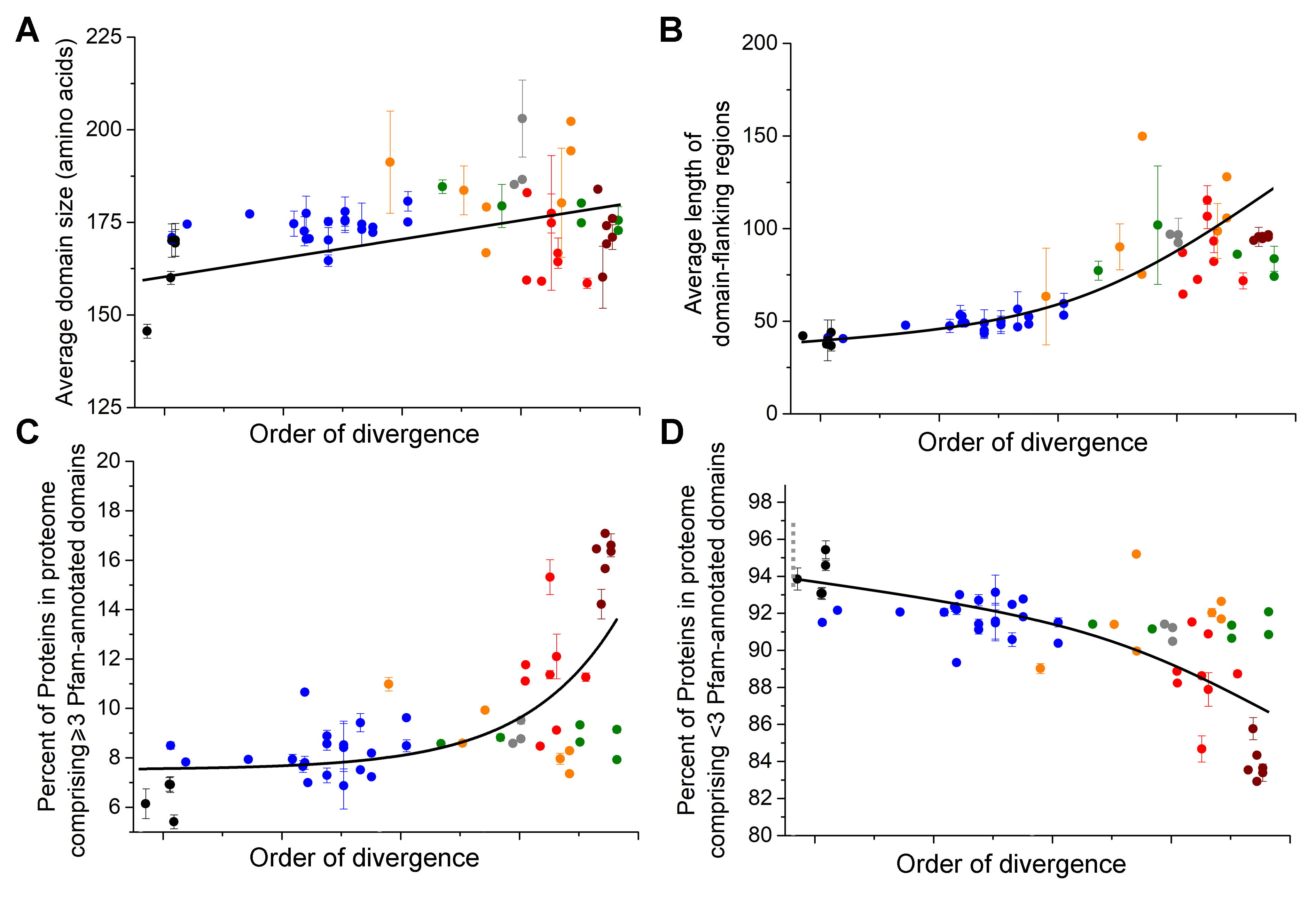

### Figure S2

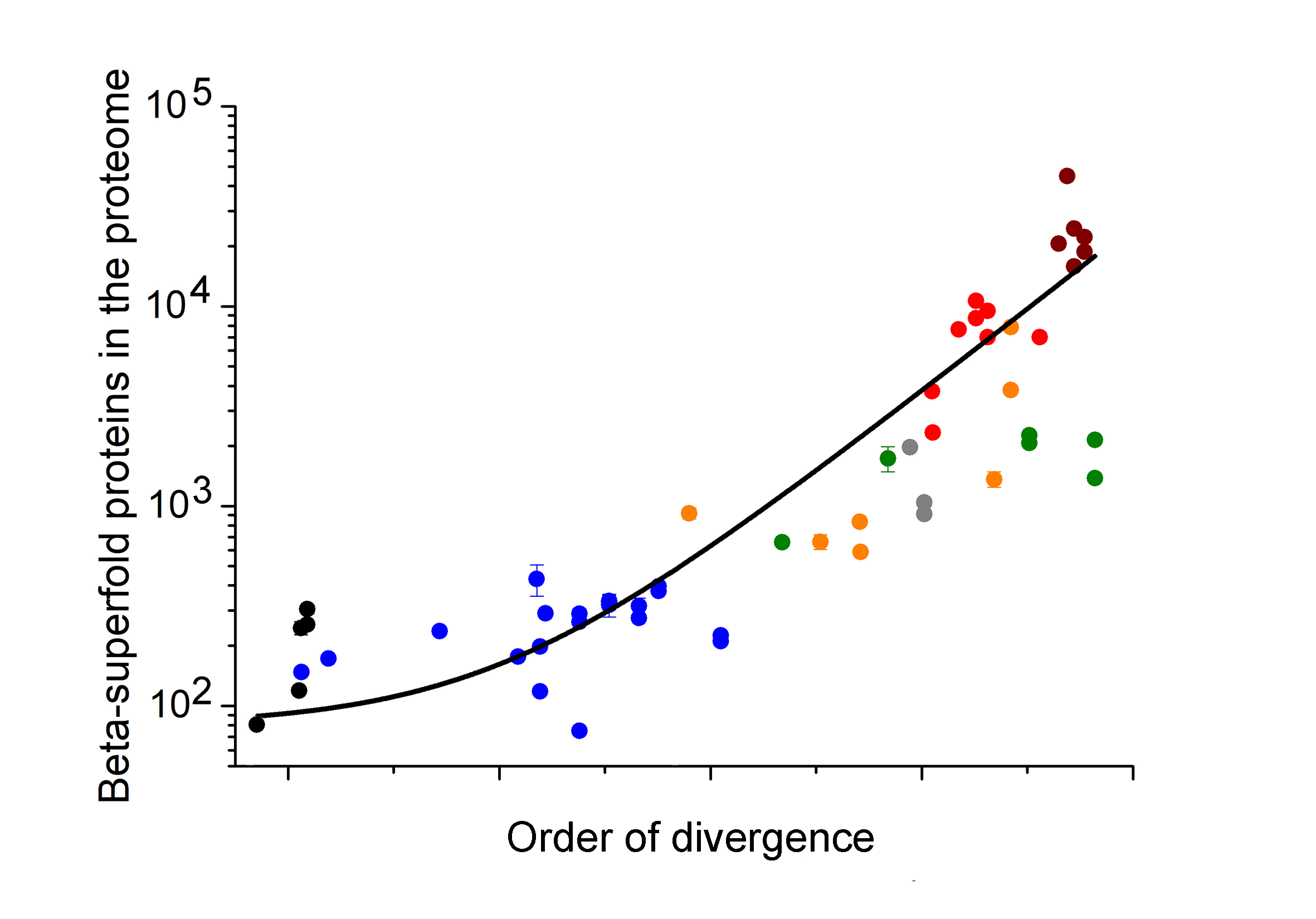

### Figure S3

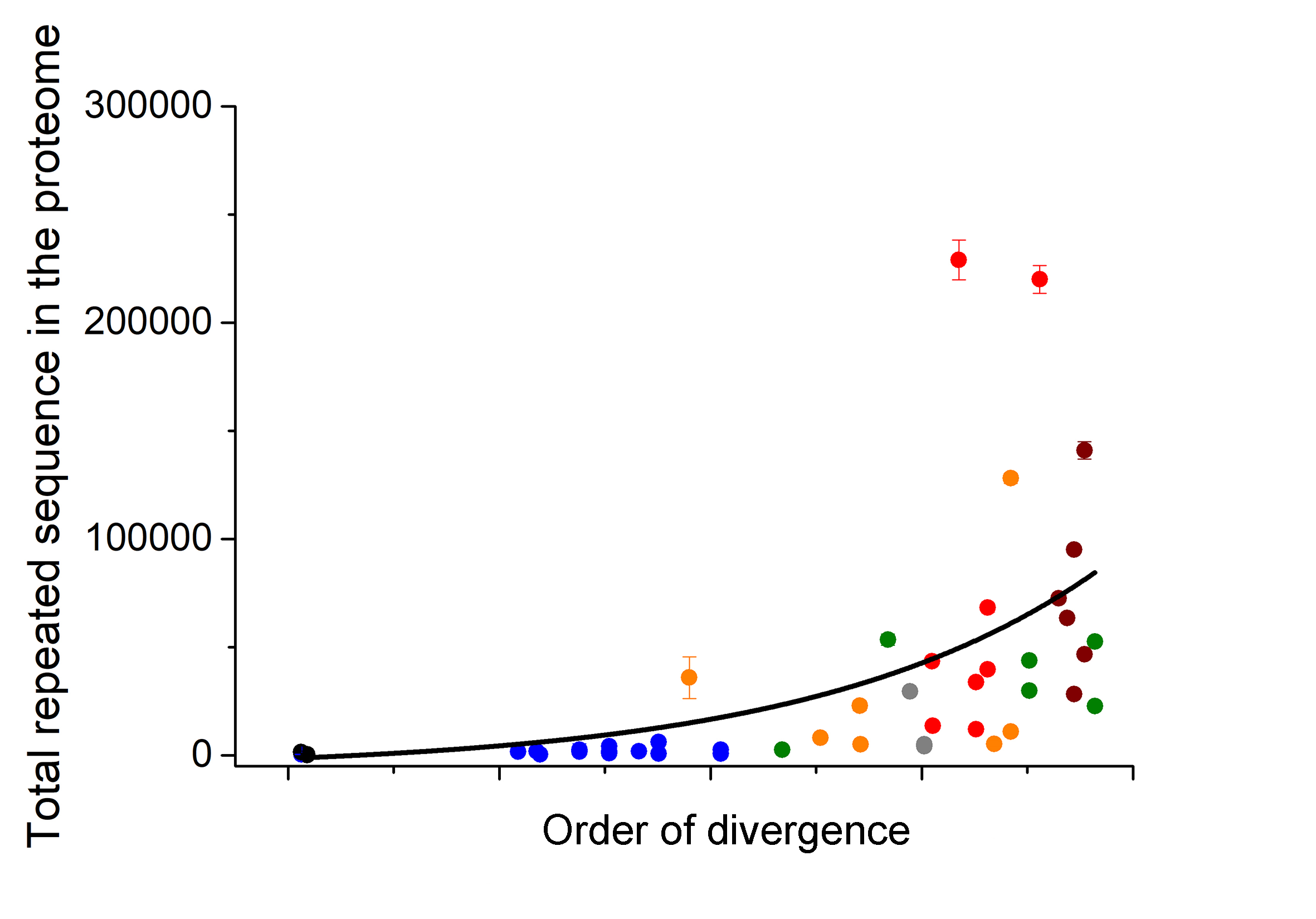

### Figure S4

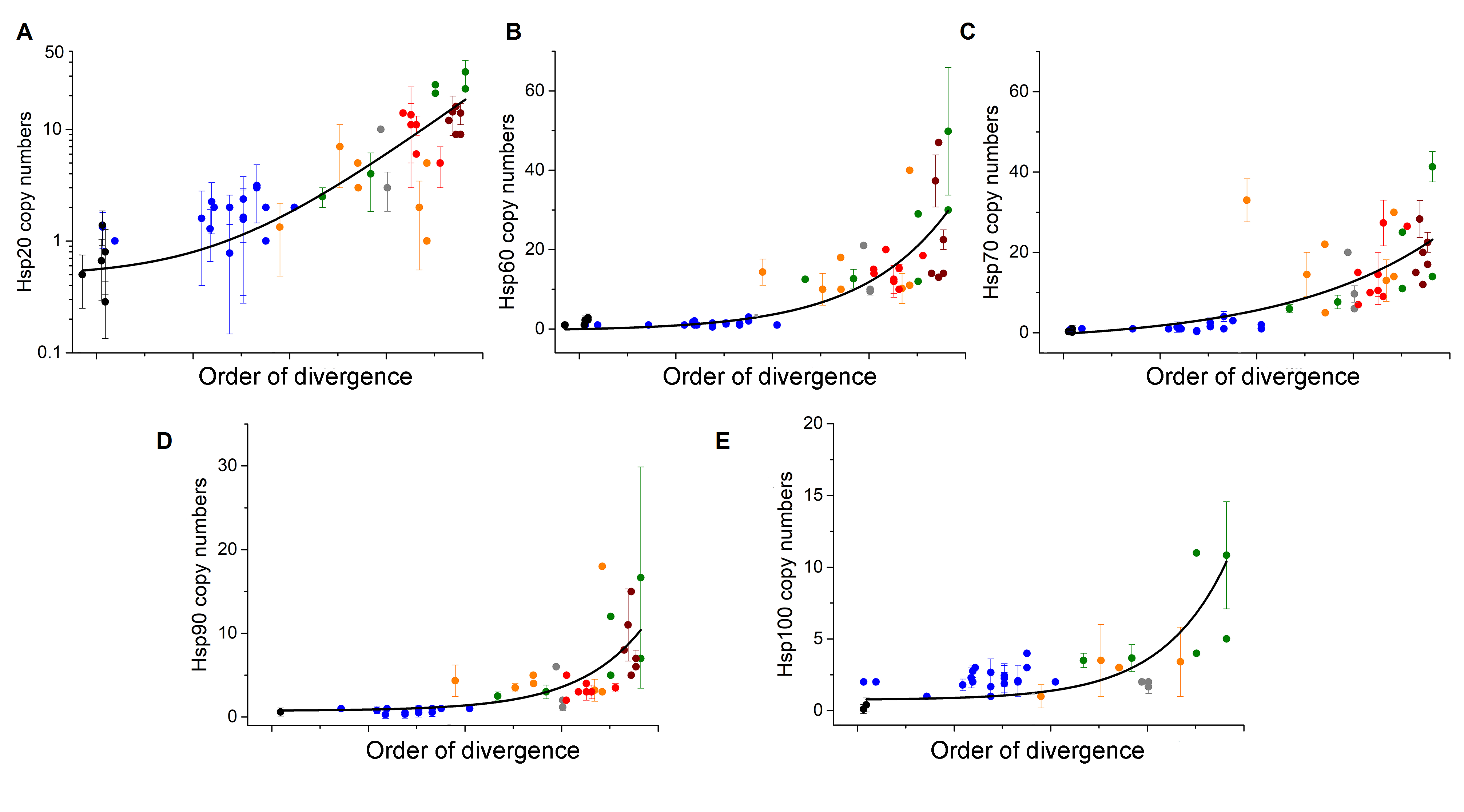
