## Supplementary material for "On the evolution of chaperones and co-chaperones and the expansion of proteomes across the Tree of Life": Response to Reviewers

**RC-2020-00333 - Point-by-Point response**

**Original comments are in blue and responses in black**

------------------------------------------------------------------------

**Reviewer #1**

**Summary**

The authors present well written work on the evolution of proteome size and complexity, and the corresponding changes in chaperone proteins. Interestingly, they find chaperone copy numbers increase linearly with proteome size, despite the increasing 'complexity' of, in particular, post-LECA genomes. They suggest that to address the rise in complexity, organisms express chaperones at higher levels and an expanding network of co-chaperones has evolved across the tree of life.

**Major comments**

**Comment-1.** Summary reads strangely relative to the rest of the manuscript, and lists facts in a way that makes the purpose of the study confusing. I think most readers will dislike the characterisation of evolution as a progress from simple to complex, and the authors' might want to avoid this language throughout the manuscript- bacteria and archaea have also been evolving over this period of times, and have not become more 'complex'? Similarly the authors should reconsider their figure legend titles. As a specific example, 'in the course of evolution' should become 'across the tree of life'.

**Response**

Thank you for these crucial suggestions. We agree with the reviewer, and with Reviewer 2 (see below) that bacteria and archaea have also been evolving since their emergence, so basically, we (humans) and the simplest archaea have the same evolutionary origin. However, we all agree that the simplest archaea/bacteria are far more similar to LUCA than we are. That said, we accept the criticism that putting our analysis in the context of evolutionary time is an over-interpretation given that we have not examined the protein/proteome phylogeny (in relation to proteome complexity; for chaperones we have). We have thus reformulated the figures and text, to a comparison across the Tree of life, rather than a time-dependent evolutionary process. Specifically: as a first step, we revised the Figures to rename the X-axis as “Order of divergence”, rather than “Divergence time (million years)” in the previous version. In the revised main text we emphasized the fact that the branch lengths of the Tree of Life represent the *relative* order of divergence of the different clades, rather than time. All instances of ‘in the course of evolution’ has been replaced by ‘across the Tree of Life’.

Secondly, we revised the main text to emphasize on prokaryote *vs.* eukaryote comparison, rather than comparing organisms that diverged at different time-points. Within bacterial and archaeal domains, proteomes do not seem to expand against the order of divergence (as the reviewer argued, bacteria and archaea have not become more complex, also see Comment-5).

Thirdly, the word ‘complexity’ has been omitted from the manuscript. The section “The expansion of proteome complexity” now reads as “Proteome expansion by *de novo* innovations”. In the previous version, *increasing complexity* in fact implied a torrent of *de novo* innovations that impose a larger burden on the chaperone machinery. Instead of ‘complexity’, the latter is clearly stated in the revised manuscript.

In the spirit of these changes, the title of the revised manuscript, figure legend titles, and related section titles have been edited as follows.

| **Submitted version** | **Revised version** |
| --- | --- |
| **Paper title**. On the evolution of chaperones and co-chaperones and the exponential expansion of proteome complexity | On the evolution of chaperones and co-chaperones and the expansion of proteomes across the Tree of Life |
| **Section title**. A Tree of Life analysis of the expansion of proteome complexity and chaperones | A Tree of Life analysis of the expansion of proteomes and chaperones |
| **Section title**. The expansion of proteome size | The expansion of proteome size across the Tree of Life |
| **Section title**. The expansion of proteome complexity | Proteome expansion by *de novo* innovations |
| **Figure 1 legend title**. Expansion of proteome size | Expansion of proteome size across the Tree of Life |
| **Figure 2 legend title**. Expansion of proteome complexity | Expansion of proteomes by *de novo* innovations |

Further, changes have been made in the Summary and in the main text to exclude any impression that proteomes/organisms have become more complex with time. Rather we emphasized prokaryote *versus* eukaryote comparison.

**Comment-2.** I think the manuscript would be improved if the authors significantly shortened the discussion of genome size evolution- this is fairly well understood, and could be covered briefly, especially as the main focus of the manuscript is on the evolution of chaperone and co-chaperone repertoire. They could also make clearer quantitative links between protein complexity and the evolution of chaperones and co-chaperones- perhaps this should be in the discussion? The authors might also consider referencing 'The evolution of genome complexity', which could be relevant to this manuscript and might make the work of broader interest.

**Response**

We thank the reviewer for this suggestion. The main focus of our paper is indeed the evolution of chaperones and co-chaperones but within the context of the expansion of proteomes. Having this focus in place, the discussion on proteome size evolution (section: The expansion of proteome size across the Tree of Life) has been revised and shortened to emphasize more on prokaryote *versus* eukaryotic comparison.

The suggestion to provide “clearer quantitative links between protein complexity and the evolution of chaperones and co-chaperones” is indeed very useful and we authors sincerely thank the reviewer. To address this suggestion we revised **Figure 4** to quantitatively compare the expansion of proteomes and that of chaperones, under one roof. This Figure compares proteome parameters that supposedly demands more chaperone action in all three domains of life and simultaneously summarizes the expansion of the chaperone machinery lacking *de novo* innovations.

The first paragraph of the Discussion section has been revised accordingly that walks the reader through the revised **Figure 4** and finally introduces to the dichotomy it implies.

We did not understand the last comment “The authors might also consider referencing 'The evolution of genome complexity', which could be relevant to this manuscript and might make the work of broader interest.” We’d be glad to address it upon further clarification.

**Comment-3.** The authors state 'protein trees were generated and compared with ToL to account for gene loss and transfer events'. The methodology for this procedure is not given in the manuscript. The authors should back up this point, and make it clear this is why they reconstruct the trees. Currently it is not convincing to me that the authors have found HGT given the considerable phylogenetic uncertainty in the basal events in the tree of life. I also expect the tree of a single protein to be potentially lack information due to the short sequence considered and possible lack of power. The authors need to consider whether the data is really of high enough quality to assess this.

**Response**

Thank you for this suggestion. For the various chaperone families, we manually compared the protein trees with the Tree of Life. This is clearly stated in the revised Methods section (see Page 25, Lines 31-32). We agree, however, that the identifying HGT, and in general, trees of single domains that are highly diverged, are tricky. We did our best to address these caveats. Specifically:

We re-evaluated our work in the light of a recent study (PMID: 32316034). This paper discussed the phylogenetic uncertainties associated with molecular dating and re-evaluated the assignment of several protein families to LUCA. A careful analysis revealed that the reviewer is indeed right, meaning many of the HGT events shown in the previous version **Figure 3B** was indistinguishable from the phylogenetic uncertainties.

Accordingly, we revised the section “The core-chaperones emerged in early-diverging prokaryotes”. We removed the previous version **Figure 3B**, along with all instances of HGT events mentioned in the main text, except one (archaea to Firmicute HGT of HSP60, which is well-supported by the data and was also detected previously). Dating the emergence of chaperone families was also re-evaluated. Though the major conclusions were not altered, we discussed the phylogenetic uncertainties associated with our work and the overall confidence of each dating analysis. We believe these discussions would be very useful to the readers.

Finally, we note that most of our key assignments (points of emergence, and major HGT events) are in agreement with previous works. Specifically: the emergence of HSP20 and HSP60 to LUCA (Sousa et al., 2016; Weiss et al., 2016) and HSP60 being horizontally transferred from archaea to Firmicute (Techtmann and Robb, 2010) and HSP20 being horizontally transferred between bacterial clades and between bacteria and archaea (Kriehuber et al., 2010).

**Comment-4.** Methods- the authors could consider taking an alternative source of LUCA proteins, rather than those found in 'Nanoarchaeota and Aquificae': it's possible these are not representative of LUCA, and it seems a somewhat arbitrary choice- the authors could consider using one of the available curated sets, such as that generated by Ranea et al. (2006).

**Response**

The reviewer is right that a more robust LUCA set could be used. However, given that the revised manuscript focuses on comparison across the ToL, and foremost on prokaryote *versus* eukaryote comparison, we don’t think that refining this set is important. Foremost, this set was used for one purpose only, for determining changes in domain length. And, the set of 38 X-groups used for this analysis are in fact, the ones present in all organisms across the ToL. Hence, we kept the original analysis, while mentioning that these 38 X-groups are conserved across the ToL, and removed the argument for LUCA assignment. See Page 5, Line 22.

**Comment-5.** The patterns observed might only hold because of differences in the taxa that diverged pre and post LECA? The authors might consider subgroup analyses to ensure this is not the case. The authors could also consider using methods that take phylogeny into account.

**Response**

The reviewer is right that within prokaryotic domains proteomes do not seem to expand. For example, excluding a few early-diverging prokaryotes and parasites, proteome size in bacteria and archaea varies within 2000-3000 proteins per proteome. Only when pre-LECA and post-LECA organisms are compared, significant differences are observed. We thank the reviewer for this suggestion. We revised the main text to focus on prokaryote *versus* eukaryote comparison. This re-focusing does not change any of our major conclusions, but rather puts our analysis in the right context (see Comment 1).

**Minor comments**

**Comment-6.** 'Life's habitability has also expanded from its 10 specific niche of emergence-likely deep-sea hydrothermal vents, to highly variable and extreme 11 ranges of temperature, pressure, exposure to high UV-light, dehydration and free oxygen.' This is not really correct, as bacteria and archaea are found worldwide, and in the most extreme environments.

**Response**

Thank you for this suggestion. We removed the above-mentioned sentence.

**Comment-7.** 'We reconciled the topology of our tree'- on first read this was not clear, I did not realise the authors were only building trees for subsets of the data- time tree is the best source for the overall topology. The phrase 'manually curated and adjusted' is used in the methods. This language is much too vague, and not a clear explanation of the steps taken.

**Response**

We apology for this confusion. The overall topology of our Tree of Life is indeed taken for TimeTree. We edited the text in Page 4, Line 4 to clarify this issue.

The obtained tree topology was manually curated and adjusted to depict eukaryotes stemming from Asgard archaea and *Alphaproteobacteria*, by an endosymbiosis event. This is clearly mentioned in the Methods section (see Page 22, Lines 24-28).

**Reviewer #2**

**Summary**

Rebeaud and colleagues analyze evolution of chaperones compared to the evolution of whole proteome complexity across the entire tree of life. Their principal conclusions are well captured in the following quote from the Discussion:

"Comparison of the expansion of proteome complexity versus that of core-chaperones presents a dichotomy-a linear expansion of core-chaperones supported an exponential expansion of proteome complexity. We propose that this dichotomy was reconciled by two features that comprise the hallmark of chaperones: the generalist nature of core-chaperones, and their ability to act in a cooperative mode alongside co-chaperones as an integrated network. Indeed, in contrast to core chaperones, there exist a consistent trend of evolutionary expansion of co-chaperones."

**Major comments**

**Comment-1.** The general theme of the evolution of proteome management is of obvious interest. Unfortunately, the entire analysis is shaky and fails to convincingly ascertain the authors' conclusions. There are many issues. Throughout the manuscript, the authors discuss 'expansion' of the proteome in bacteria, archaea and eukaryotes, creating the impression of a consistent evolutionary trend. No such trend actually exists if one considers the means or medians of proteome sizes within each of the three domains of life (there is a transition to greater complexity in eukaryotes). The maximum complexity, certainly, increases with time which can be attributed to the 'drunkard's walk' effect. This hardly qualifies as 'expansion'.

**Response**

The reviewer is right that within prokaryotes proteomes do not seem to significantly expand. Reviewer-1 raised a similar concern that prokaryotes and eukaryotes have been evolving for the same period of time and have not expanded significantly. We understand the misconception instated by the earlier version and we thank the reviewers for pointing it out. Accordingly, we revised the main text to clarify these issues, as described in the following.

Firstly, the main text was revised to emphasize on prokaryote *versus* eukaryote comparison. The reviewer agrees that compared to prokaryotes, “there is a transition to greater complexity in eukaryotes”. This re-focusing does not change any of our major conclusions, but rather provides a systematic comparison that is adequately supported by data.

Secondly, we revised the Figures to rename the X-axis as “Order of divergence”, rather than “Divergence time (million years)” in the previous version. We emphasized the fact that the X-axis actually represent the *relative* order of divergence of the different clades, rather than absolute dates. This emphasis certainly does not create the impression of a consistent evolutionary trend. Instead, combined with the revised main text, it depicts that only when pre-LECA and post-LECA organisms are compared, clear trends of proteome expansion is observed.

**Comment-2.** The authors further claim a 'linear' expansion of the chaperone set and 'exponential' expansion of the total proteome size. These are precise mathematical terms and, as such, require fitting to the respective functions. No such thing in this manuscript. Even apart from that shortcoming, the explanation of both 'linear' and 'exponential' are quite confusing. Thus, when explaining the 'linearity' of chaperone evolution, the authors refer to the lack of major innovation among the chaperones. This is correct in itself but has nothing to do with linearity. Apart from the aforementioned conceptual problems, the estimation of the 'exponential' growth of the proteome are naive, inconsistent and inaccurate.

**Response**

Our uses of ‘linear expansion’ versus ‘exponential expansion’ may have been confusing although we have defined quite clearly what we mean by that (*i.e.*, that it is not the mathematical sense). The statement regarding “the lack of major innovation among the chaperones” was made in this context/definition and was consistent with it.

Nonetheless, to avoid confusion, we revised the main text by excluding the ‘linear expansion’ and ‘exponential expansion’ terms. We simply stated that a torrent of *de novo* innovations has occurred during the expansion of proteomes from prokaryotes to eukaryotes. In contrast, the evolutionary history of core-chaperones lacks such major innovations. Accordingly, the title of the revised manuscript, figure legend titles, and related section titles have been edited as follows.

| **Submitted version** | **Revised version** |
| --- | --- |
| **Paper title**. On the evolution of chaperones and co-chaperones and the exponential expansion of proteome complexity | On the evolution of chaperones and co-chaperones and the expansion of proteomes across the Tree of Life |
| **Section title**. A Tree of Life analysis of the expansion of proteome complexity and chaperones | A Tree of Life analysis of the expansion of proteomes and chaperones |
| **Section title**. The expansion of proteome complexity | Proteome expansion by *de novo* innovations |
| **Figure 1 legend title**. Expansion of proteome size | Expansion of proteome size across the Tree of Life |
| **Figure 2 legend title**. Expansion of proteome complexity | Expansion of proteomes by *de novo* innovations |

**Comment-3.** As the base point for the expansion estimates for archaea and eukaryotes, the authors take parasitic forms. Even leaving aside the highly dubious claims that these organisms belong to the clades that diverged first from the respective ancestors, parasites are not an appropriate choice for such estimates because they certainly are products of reductive evolution. For bacteria, inconsistently, the authors choose a free-living form from a dubious ancient clade, and not even the one with the smallest genome. All taken together, this robs the expansion estimates of any substantial meaning.

**Response**

This point is overall valid. Although we adamantly reject the insinuation of “dubious claims that these organisms belong to the clades that diverged first from the respective ancestors” – firstly, we did not make any claims to this end, but took the ToL constructed by others (Hedges et al., 2015); second, that these claims are dubious need to backup by counter-evidence/data and with all due respect, neither were provided by the reviewer. However, what is of concern is that in a symbiont/parasite chaperones of the host may have a key role, and thus the comparison to free-living organisms could be misleading. To address this concern we excluded the obligatory endosymbiont *Nanoarchaeum equitans* and the parasitic organisms from the expansion estimates and such discussions are now limited to free-living organisms only. Further, as described in response to Comment-1, the revised manuscript focuses on prokaryote versus eukaryote comparison.

Note that phylogenetic analysis often assigns parasitic and symbiotic organisms that have experienced reductive evolution as the earliest diverging clades of their corresponding kingdoms of life. Examples include *Nanoarchaeum equitans*, an obligate symbiont, assigned as the earliest diverging archaea (Hedges et al., 2015; Huber et al., 2002; Waters et al., 2003), and parasitic Excavate assigned as one of the earliest diverging eukaryotes (Burki et al., 2020; Simpson et al., 2002). In accordance with these studies, these parasitic and symbiotic organisms were included in our analysis. We acknowledged this fact in the Methods section (see Page 22, Lines 9-16).

**Comment-4.** The authors do make a salient and I think essentially correct observation: chaperones typically comprise about 0.3% of the proteins in any organism. As such, this presents no dichotomy in evolutionary trends to be explained. Surely, as examined and discussed in the paper, eukaryotes also show significant increases in the size and domain content of the encoded proteins, suggesting the possibility that might need more chaperones. However, if this is the explanandum, rather than the number of proteins in the proteome as such, it should be clearly stated. Furthermore, it is quite natural to assume that this increase in protein complexity without a commensurate increase in the chaperone diversity, is enabled by higher expression of the chaperones as suggested in the Discussion of this paper. I doubt there is any big surprise here and even much need for an extended discussion let alone a special publication.

**Response**

As emphasized, and shown, eukaryotes have not only larger proteomes in terms of the number of proteins or protein size. They have a higher content of proteins that are prone to misfolding. This is shown explicitly, in Figure 2 (namely, multidomain proteins, repeat, beta-rich proteins, etc’) and is reiterated in a summary figure (suggested by Reviewer 1). Further, in response to Reviewer-3’s suggestion, we showed that eukaryotes feature much higher proportions of aggregation-prone proteins per proteome than prokaryotes (**Figure 2E**).

To further clarify, we revised **Figure 4** to quantitatively compare the expansion of proteomes and that of chaperones, under one roof. This Figure compares proteome parameters that supposedly demands more chaperone action in all three domains of life and simultaneously summarizes the expansion of the chaperone machinery lacking *de novo* innovations.

In addition, the first paragraph of this Discussions section is revised to state that from prokaryotes to eukaryotes, proteomes have expanded by duplication-divergence as well as by innovations (*de novo* emergence of new folds). Thus, it’s not about the size only (a challenge that a proportion expansion of chaperone genes would resolve, *i.e.*, the 0.3%) but about proteome composition changing in a way that demands more and more chaperone action.

We also agree with the assertion that “it is quite natural to assume that this increase in protein complexity without a commensurate increase in the chaperone diversity, is enabled by higher expression of the chaperones”. However, we belong to a group of scientists for whom natural assumptions are insufficient, and think that supporting evidence is of importance.

**Reviewer’s significance statement**

As such, in the opinion of this reviewer, there is no substantial advance over the existing knowledge in this paper. Should the authors wish to revise, they would need to develop robust methodology to measure proteome expansion. That would involve starting from reconstructed ancestors rather than any extant forms (let alone parasites). I doubt that such analysis, non-trivial in itself, reveals an strong, consistent trends other than the well known increase in complexity in eukaryotes.

**Response**

We agree that to assert evolutionary, time-dependent trends one needs to analyze phylogenies and reconstructed ancestors, but still think that a comparison of proteome and chaperone contents along the Tree of Life is meaningful. We thus respectfully, yet adamantly disagree with “no substantial advance over the existing knowledge”. We strongly believe, as does Reviewer-3, that the results and the model presented in this paper are “fascinating to consider and… will stimulate a good deal of important discussion…”.

**Reviewer #3**

**Summary**

The manuscript by Rebeaud et al describes phylogenetic analyses of proteome and chaperone complexity. The authors analyzed species across the tree of life to predict the proteome and chaperone properties of ancestors spanning to the last universal common ancestor. Their analyses indicate that many proteome properties increased in complexity over evolutionary time including: average protein length, the number of multi-domain proteins, the size of the proteome, the number of repeat proteins, and the number of beta-superfold proteins that are known to be difficult to fold. Their analyses also indicate an expansion in chaperone families that corresponds to the increase in proteome complexity. Based on their analyses, the authors propose a model where early life relied on a limited number of chaperones (Hsp20 and Hsp60) and that as proteome complexity evolved, so did chaperone complexity. Core chaperones including Hsp90, Hsp70, and Hsp100 evolved relatively early, and later chaperone evolution was driven by the appearance and alterations of co-chaperones and auxiliary factors as well as by increases in the protein abundance of chaperones.

**Major concerns**

**Comment-1.** This work is appropriately based on phylogenetic inferences, but as such, the limitations and uncertainties of phylogenetic inferences need to be discussed. This in no way takes away from the work, quite the opposite, it would make it richer by encouraging broader interpretations where justified and clear understanding of where support for the model is strongest. Posterior probabilities need to be discussed and the range of properties that a likely ancestor might have based on the data should be discussed. How this impacts the conclusions and models should be discussed. Throughout the manuscript, the authors present most-likely ancestral models (as I understood it), what are the next most likely models? How much power is there to distinguish one model from another? It would be very helpful to have a section describing the limitations and uncertainties of the phylogenetic analyses and how these relate to the main findings and conclusions.

**Response**

We thank the reviewer for this suggestion. Reviewer-1 raised a similar suggestion (see Comment-3). The phylogenetic analysis in our paper included dating the emergence of core- and co-chaperone families, and attempt to infer major their HGT events, foremost in relation to the origin of eukaryotic chaperones. To highlight the uncertainties of phylogenetic inferences we re-evaluated our work in the light of a recent study (PMID: 32316034) that carefully analyzed the uncertainties associated with the assignment of several protein families to LUCA.

Ideally, for a protein family to be assigned to LUCA, there must be a single split of bacterial and archaeal domains at the root of the protein tree with strong bootstrap support, and the inter-domain branches would be longer than the intra-domain branches (PMID: 32316034). In the revised main text we discussed that only the HSP60 protein tree satisfies this criterion. HSP20 protein tree depicts a clear single split of bacterial and archaeal domains at the root, albeit with weak bootstrap support, and inter-domain branch lengths are smaller than intra-domain branch-lengths. We discussed that this is indeed the case of phylogenetic uncertainty, which means the sequence of this small, single-domain chaperone lacks the information to make reliable inference at the basal events in the ToL.

In addition, the HGT events discussed in the previous version appear to be indistinguishable from phylogenetic uncertainties and we removed all instances of HGT events mentioned in the main text as well as Figure 3B. Only one HGT event – HSP60 being horizontally transferred from archaea to Firmicute, which is well-supported by the data is kept in the revised main text. We believe these discussions would be very useful to the readers.

Finally, we note that most of our key assignments (points of emergence, and major HGT events) are in agreement with previous works. Specifically: the emergence of HSP20 and HSP60 to LUCA (Sousa et al., 2016; Weiss et al., 2016) and HSP60 being horizontally transferred from archaea to Firmicute (Techtmann and Robb, 2010) and HSP20 being horizontally transferred between bacterial clades and between bacteria and archaea (Kriehuber et al., 2010).

**Comment-2.** General features that impact foldability, including contact order, should be discussed and what features can be searched for in genomes that relate to these - e.g. beta-rich proteins.

**Response**

Thanks for this valuable idea! Contact order, and other predictors of problematic folding are highly relevant but their analysis is structure-based and hence inapplicable on the proteome (sequence) scale. We did, however, estimate the proportion of aggregation-prone proteins in the proteome. These proteins were identified by CamSol method that assigns poorly soluble regions from sequence data. Indeed, some of these predicted ‘poorly soluble segments’ refer to the hydrophobic core of the respective folded state instead of ‘true’ aggregation hotspots. With this unavoidable potential caveat, it appears that compared to prokaryotes, aggregation-prone proteins in the proteome have become nearly 6-fold more frequent in Chordates.

Following changes were made to accommodate this new analysis:

Figure 2 is revised to include a new panel (panel-E) that shows the expansion of aggregation-prone proteins in the proteome across the Tree of Life. The same result is summarized in the summary Figure 4.

A new paragraph entitled “*Proteins predicted as aggregation-prone became ~6-fold more frequent in the proteome*” is added to the Results section, which describes the principle and the main results (see Page 7, Lines 14-28).

The methodology is included in the Methods section, in a paragraph entitled “*Predicted proportion of aggregation-prone proteins in the proteome*”, see Page 24 Lines 17-27. For each representative organism, the percent of aggregation-prone proteins in proteome data are provided as Data S10.

This analysis is also included in the revised Abstract: “Proteins prone to misfolding and aggregation, such as repeat and beta-rich proteins, proliferated ~600-fold, and accordingly, proteins predicted as aggregation-prone became 6-fold more frequent in mammalian compared to bacterial proteomes.” See Page 2, Lines 7-9.

**Comment-3.** "Core" chaperones needs to be defined.

**Response**

Thank you for this suggestion. We restructured Page 3 Lines 19-23 in the Introduction to clearly explain this aspect. The current text is quoted below.

“Chaperones can be broadly divided into core- and co-chaperones. Core-chaperones can function on their own, and include ATPases HSP60, HSP70, HSP100, and HSP90 and the ATP-independent HSP20. The basal protein holding, unfolding, and refolding activities of the core-chaperones are facilitated and modulated by a range of co-chaperones such as J-domain proteins (Caplan, 2003; Duncan et al., 2015; Schopf et al., 2017).”

**Minor concerns and thoughts**

**Comment-4.** This manuscript stimulated me to think about the dynamics between chaperone evolution and proteome evolution. The ability to tolerate proteins that need chaperones seems linked to major evolutionary innovations. Once you have these innovations though, you are addicted to the chaperones - and an expansion of the number of sub-optimal proteins. These ideas seem like they would be valuable to include in the discussion of this work. More generally, it would be wonderful to have a discussion of future directions that this work may spark.

**Response**

This is indeed a fascinating question or set of questions, that we have also become intrigued about following this work, We introduced a short section, though more of an ‘appetizer’ than a detailed discussion, as we know almost nothing about the co-evolution of new proteins and chaperones.

**Reviewer’s significance statement**

This manuscript provides a fascinating glimpse back in time of a fundamental interplay - between chaperone evolution/addiction and proteome evolution. I am not an expert in phylogenetic analyses so I cannot judge the details of the analyses. As an expert in molecular evolution and chaperones, I found the approach and model fascinating to consider and I believe it will stimulate a good deal of important discussion in these fields. I have one major concern that I feel ought to be addressed in the manuscript and a number of points that I would encourage the authors to consider. I am sure that these can be readily addressed and I look forward to seeing this work published and the further discussion and ideas that it may stimulate.

**Response**

Thank you!
